# Phylogenomics and biogeography of the parrot genus *Pyrrhura* with implications for systematics and conservation

**DOI:** 10.64898/2026.01.21.700815

**Authors:** Jaime G. Morin-Lagos, Taylor Hains, José Cerca, Michael Wink, Stacy Pirro, Cristina Y. Miyaki, Shannon J. Hackett, John M. Bates, Michael D. Martin

## Abstract

The genus *Pyrrhura* (Psittacidae: Arini) is one of the most diverse groups of Neotropical parrots. Its species are charismatic, widely kept as pets, and frequently bred outside their native ranges. Yet, nearly half are currently listed as threatened by the IUCN within their natural distributions. Conservation assessments and population estimates often depend on the validity of accepted taxonomic boundaries. However, despite previous systematic efforts, the evolutionary relationships among and within many *Pyrrhura* species remain poorly resolved, largely due to a recent and rapid radiation. Here, we generated whole-genome sequences for all currently recognized *Pyrrhura* species, including multiple intraspecific taxa, to reconstruct a robust nuclear phylogeny under the multi-species coalescent model, alongside the most comprehensive mitogenome-based phylogeny of the genus to date. Although both phylogenies supported the monophyly of most currently accepted species, we identified several instances of mito-nuclear discordance, particularly involving the placement of early-diverging lineages, which are best explained by incomplete lineage sorting and historical gene flow. Additionally, we detected three distinct captive lineages that do not cluster with any known wild populations, suggesting substantial overlooked genetic diversity in the world’s captive populations. Ancestral range reconstructions indicate multiple and relatively recent colonization events into the northern and central Andes, likely associated with the uplift of the Andes and the emergence of new ecological niches. Together, our results reveal a complex evolutionary history in *Pyrrhura*, shaped by rapid radiations, incomplete lineage sorting, and gene flow. We show that integrating nuclear and mitochondrial data with broad geographic and taxonomic sampling, including captive individuals, can uncover overlooked genetic diversity and help to resolve long-standing systematic uncertainties. Finally, we show that several topological discrepancies among previous studies can be attributed to differences in sampling strategies, particularly within the most polytypic *Pyrrhura* species.

## Introduction

*Pyrrhura* Bonaparte (1856) is the second most species-rich genus of Neotropical parrots (Arinae), distributed from southern Central America through tropical and subtropical South America (Gill et al., 2025). The genus currently comprises 24 currently recognized species, 11 of which are polytypic, reflecting substantial geographic and phenotypic variation. This diversity, together with a broad ecological and biogeographic range, makes *Pyrrhura* an important system for studying avian diversification in the Neotropics.

Many *Pyrrhura* species are charismatic and widely maintained in captivity, having been exported and bred outside their native ranges for multiple generations, particularly in Europe and North America following their early popularity prior to the establishment of modern international biodiversity regulations such as CITES (1975) and the Nagoya Protocol on Access and Benefit Sharing (ABS, 2010). In contrast, wild populations across the Neotropics have faced ongoing threats from habitat loss, fragmentation, climate change, and illegal capture. As a result, 15 *Pyrrhura* species are currently listed as Vulnerable, Endangered, or Critically Endangered by the IUCN (2025). This disparity between captive abundance and wild decline raises concerns about the erosion of genetic diversity in natural populations and the potential loss of unique evolutionary lineages.

Effective conservation planning depends on robust taxonomic frameworks, yet species delimitation and phylogenetic relationships within *Pyrrhura* have long been considered problematic. Morphological differentiation among some taxa is subtle, while traits such as plumage coloration can be highly variable within and among populations, complicating species diagnoses. Consequently, the taxonomy of the genus has undergone multiple revisions as molecular data have become available. The first comprehensive molecular phylogeny of *Pyrrhura* by Ribas et al. (2006) identified three major evolutionary lineages: *P. cruentata* from the Atlantic forest, the *P. picta-leucotis* complex, and a third clade comprising the remaining species. The latter two distributed across the humid Neotropical forests.

Further molecular examination of the *P. picta*-*leucotis* complex revealed six subclades in the group (Arndt & Wink, 2017). However, this and other phylogenetic efforts (Provost et al., 2018; Wright et al., 2008) relied on a limited number of mitochondrial and nuclear loci and included incomplete taxon sampling. Such datasets are often insufficient to resolve relationships within lineages shaped by recent and rapid diversification, a pattern inferred for *Pyrrhura* (Ribas et al., 2006). More recently, a phylogeny based on ultraconserved elements (UCEs) included nearly all IOC-recognized *Pyrrhura* species, except for *P. pfrimeri* (Smith et al., 2023), but still lacked complete subspecies representation, underscoring persistent gaps in genus-wide resolution.

Considering all IOC-recognized subspecies, *Pyrrhura* comprises 48 taxa, most of which have never been included in molecular phylogenetic analyses due to their uncertain systematic status. Several subspecies have been proposed for elevation to species-rank, such as *P. p. subandina* (Todd, 1917), while new taxa, e.g. *P. amazonum araguaiaensis* (Arndt & Wink, 2017), have been described based on morphological differentiation and geographic isolation. However, many of these proposals have not been adopted by major taxonomic authorities, including the IOC World Bird List and the South American Classification Committee (SACC) (2025), largely due to limited or inconclusive molecular evidence, as illustrated by the SACC Proposal 524 (Penhallurick, 2012). As a result, the evolutionary relationships of several lineages, including species complexes such as *P. melanura* (Ridgely & Robbins, 1988), remain unresolved, hindering both evolutionary inference and the identification of appropriate conservation units.

Integrating mitochondrial and nuclear genomic data offers a powerful framework to study complex evolutionary histories, particularly in groups where incomplete lineage sorting and interspecific gene flow may have played important roles, and to identify interspecific hybridization, which appears to be common in *Pyrrhura* (Somenzari & Silveira, 2015; Urantowka et al., 2016). Here, we present the first genus-wide phylogenomic analysis of *Pyrrhura*, integrating complete mitochondrial genomes with nuclear genomic data to infer phylogenetic relationships, estimate divergence times, assess mito-nuclear discordance, and reconstruct biogeographic history across all currently IOC-recognized species. This study provides the most comprehensive and well-supported phylogenetic hypothesis for the genus to date, reveals overlooked genetic diversity, and offers a robust genomic framework to inform future systematic revisions and conservation priorities.

## Material and Methods

### Taxon sampling

We obtained samples representing the 24 *Pyrrhura* species currently accepted by the IOC World Bird List v15.1 (Gill et al., 2025). To account for the substantial and potentially cryptic diversity within the genus, we also included nine additional taxa recognized at the subspecies level by the IOC. In addition, we incorporated DNA samples from Arndt and Wink (2017) representing three subspecies proposed by the authors, *Pyrrhura amazonum araguaiensis*, *P. lucianii orosaensis*, *P. (roseifrons) dilutissima pareneensis*, as well as the earlier proposed taxon *P. amazonum microtera* (Todd, 1947). Morphological identifications for these specimens follow the designations provided in the original study. An overview of the accepted and additionally proposed *Pyrrhura* taxa and their sampling for this study can be found in Table S1.

A total of 78 *Pyrrhura* samples representing 32 taxa were sourced from various natural history collections and from the DNA bank of the Institute of Pharmacy and Molecular Biology (IPMB at Heidelberg, Germany; now housed at Senckenberg Naturhistorische Sammlungen Dresden, Senckenberg, Leibniz Institution for Biodiversity and Earth System Research, Dresden, Germany), which holds a large collection of morphologically curated captive parrot samples (Arndt & Wink, 2017), but also wild specimens. We also included six Neotropical parrot species as outgroups. For nuclear genomic analyses, we prioritized samples originating from native geographic ranges, resulting in 33 *Pyrrhura* samples representing the 24 accepted species and eight subspecies. Details on sample sources, sequencing statistics, and accession numbers are provided in Table S2.

### DNA extraction

DNA extracts employed in the cyt-b based phylogeny (Arndt & Wink, 2017) were provided by the Institute of Plant Molecular Biology (IPMB). In addition, toe pad and tissue samples were obtained from museum collections. DNA extraction of samples sourced from museums in the United States were processed as in Hains et al. (2022). It is important to note that most of the genomic data produced at Field Museum have been previously released by the authors, although they have not yet been part of any evolutionary analysis. The rest of the samples were processed at the Norwegian University of Science and Technology (NTNU) utilizing the DNeasy Blood and Tissue kit (Qiagen Inc., Valencia, CA, USA), according to the manufacturer’s instructions.

### Genomic library preparation and sequencing

Genomic DNA samples extracted at IPMB and at NTNU-VM molecular laboratory were built into libraries at NTNU-VM. Genomic DNA was previously sheared using Covaris ME220 ultra-sonicator to a mean fragment length of 350 bp. Double-strand DNA library preparation followed the BEST protocol (Carøe et al., 2018), in which custom BGI adapters were ligated to the DNA fragments. Indexing PCR was performed using custom index primers to obtain dual-index libraries. Indexing PCR was carried out using 10 μl of library template, AmpliTaq Gold DNA polymerase (0.05 U/μl), 1X AmpliTaq Gold buffer, 0.2 mM each deoxynucleotide triphosphate (dNTP), 2.5 mM MgCl_2_, bovine serum albumin (0.4 mg/ml), 0.2 μM sample-specific forward index primer, 0.2 μM sample-specific reverse index primer, and molecular-grade water to obtain a 100-μl reaction volume. The optimal number of PCR cycles was selected for each sample based on quantitative PCR. Library amplification was carried out using the following cycling conditions: An initial step of 95°C for 10 min, then a library-specific number of cycles of 95°C for 30 s, 60°C for 1 min, and 72°C for 45 s, and a final extension of 72°C for 5 min. Amplified indexed libraries were purified with solid-phase reversible immobilization (SPRI) beads (Rohland & Reich, 2012) and eluted in 33 μl of Qiagen EB buffer. Fragment length, distribution, and molarity of the indexed libraries were assessed using an Agilent TapeStation 4200 with a high sensitivity D1000 ScreenTape (Agilent). Libraries were pooled based on these values. Low-coverage paired-end 150-bp format sequencing was performed either at the BGI Europe sequencing facility on the DNAseq platform or at the Novogene Europe sequencing facility on the Illumina HiSeq 2500 platform.

A subset of samples, including all those extracted at the Field Museum, was sequenced to higher depth (Hains et al., 2022) (Table S2). This choice was based on the availability of voucher specimens to confirm their taxonomy. These samples are primarily held in museum collections and were collected in the wild. DNA extracts from these specimens were prepared for sequencing using the Illumina TruSeq kit following the manufacturer’s instructions. The libraries were then sequenced on an Illumina HiSeq platform in paired-end 150-bp format.

### Mitochondrial genome analysis

#### Bioinformatic processing, de-novo assembly, and annotation

The bioinformatics pipeline paleomix v1.2.13.8 (Schubert et al., 2014) was used to process the raw sequencing reads. Adapter sequences and low-quality bases were trimmed using AdapterRemoval v2.3.1 (Schubert et al., 2016). Mitochondrial genomes were assembled *de novo* using NOVOPlasty v4.3 (Dierckxsens et al., 2017) and SPAdes v3.13 (Bankevich et al., 2012). Typically, both methods retrieved complete mitochondrial genomes identical to each other; other times, NOVOPlasty could not retrieve the full sequence length. However, it was possible to manually merge the contigs obtained using SPAdes to obtain full mitochondrial sequences. Initial annotation of one mitogenome sequence (*P. r. parvifrons* MUSM 32747) was carried out using the MITOS2 tool within the galaxy platform (https://usegalaxy.eu/). Additional search of tRNA genes was performed using tRNAscan-SE 2.0 (Lowe & Chan, 2016), see supplementary materials. Annotations of the protein-coding genes were refined by alignment using the Geneious Prime 2024.0.7 platform following the *Pyrrhura rupicola* NC_028404 annotation (Urantowka et al., 2016) as a reference. Then we used the transfer annotations tool (implemented in Geneious Prime) to annotate the remaining mitogenomes. We recovered 75 full and three partial mitochondrial genomes from the 78 *Pyrrhura* individuals. For the mitochondrial analysis, we included all the samples sourced. In addition we recovered full mitogenomes of the six outgroup samples. The sequences were deposited in GenBank (NCBI) under the accession codes PV917592-PV917648.

#### Sequence alignment and maximum-likelihood phylogenetic analysis

The assembled mitogenome sequences were aligned using MUSCLE v3.8 (Edgar, 2004) and manually checked on Geneious Prime. We inferred a maximum-likelihood tree using IQ-TREE v2.0.3 (Minh et al., 2020) with 1000 rapid bootstraps and using the “-m MFP+MERGE” option. This option performs nucleotide substitution model selection and chooses the best-fit partitioning scheme utilizing an implementation of PartitionFinder (Lanfear et al., 2012). The partition scheme obtained was also implemented in the BEAST analysis and can be found in Table S6. In addition, we evaluated the genetic distances between species and putative taxa using the Species Delimitation tool (Masters et al., 2011) implemented in Geneious Prime.

#### Time-calibration and biogeographic analysis

We inferred a Bayesian tree and estimated divergence times using BEAST2 v.2.7.6 (Bouckaert et al., 2019), employing the four partitions previously identified by IQ-TREE. We set all the parameters and created a .xml file using BEAUTi v2.7.6. We linked clock and tree models and maintained site models as unlinked. Substitution models for each partition were estimated using the BEAST Model Test, and base frequencies and mutation rate were set to “Empirical”. A relaxed Clock Log-Normal model was assumed, and the Yule process of speciation was set as the tree prior. Due to a lack of fossil records from closely related lineages, we opted for two secondary calibration priors extracted from the fossil-based Psittaciformes time-tree estimated by Smith et al. (2023). We employed specifically the crown age of the *Pyrrhura* genus estimated with 95% HPD covering 2.22-9.9 Mya (lognormal: M = 5.145, S = 0.4), and the divergence of *Pyrrhura* from the rest of the Arini tribe, represented here by *Ara*, *Aratinga*, *Psittacara*, and *Anodorhynchus*, estimated with 95% HPD covering 6.24-16.94 Mya (lognormal: M = 10.472, S = 0.255). We then executed two MCMC chains (each with 100,000,000 generations sampled every 10,000 generations). We checked convergence, posterior distribution, and effective sample size (ESS > 200) using Tracer v1.7.2 (Rambaut et al., 2018) and discarding the first 10% of generations as burn-in. The two independent runs were combined using LogCombiner v2.7, and a maximum clade credibility tree was created using TreeAnnotator v2.7.6.

#### Historical biogeography

We utilized the R package BioGeoBEARS (Matzke, 2013) using the script provided by the author as baseline (http://phylo.wikidot.com/biogeobears#toc19) to infer the ancestral geographical ranges of the genus *Pyrrhura* based on the time-calibrated tree. We categorized each species into five discrete areas representing different South American biogeographical regions based on Morrone et al. (2022) and the distribution of *Pyrrhura* species. These regions were defined as follows: North South America and Meso America (NSM), North Amazonia (NAM), South Amazonia (SAM), Dry Diagonal (DDI), and Atlantic Forest (ATF). Area assignment and details on the provinces included within those areas are in Table S7. Prior to running BioGeoBEARS, we used the iTOL ‘delete leaf’ function to trim the mitogenome phylogeny to include only one representative sample per species, excluding captive clades (defined as monophyletic clades represented solely by captive individuals) and divergent captive specimens. We evaluated the three models implemented in BioGeoBEARS, DEC, DIVALIKE, and BAYERALIKE, which have two free parameters for dispersal (“d”) and extinction (“e”). In addition, we tested the effect of adding the founder-effect (“j”) parameter to those three models, resulting in tests of six models in total. Given that the range of all the living species falls mainly within a single defined geographical area, which is extensive in area, we decided to set the “max_range_size” parameter to 2, indicating that ancestral species can only occur in a maximum of 2 of the 5 defined geographic areas. We then followed the Akaike information criterion (AIC) to select the likelihood version of the dispersal–vicariance analysis model with jump dispersal (DIVALIKE+J) as the best fitting model.

### Nuclear genome analysis

We used PALEOMIX pipeline v1.2.13 (Schubert et al., 2014) to process the raw sequence data. We first performed adapter trimming and paired-read collapsing for overlaps of at least 11 bp using AdapterRemoval v2.3.1 (Schubert et al., 2016). Cleaned reads ≥ 25 bp in length were mapped to a scaffold-level reference genome for *Pyrrhura perlata* (FMNH voucher: 389699; Hains et al. *in prep*) using BWA-mem v0.7.15 (Li & Durbin, 2009) and default settings, keeping only those aligned reads with a Phred-scaled mapping quality (MAPQ) ≥ 30. We estimated endogenous DNA content, sequencing depth, and clonality using the PALEOMIX summary files.

We conducted a preliminary assessment of the nuclear data for all *Pyrrhura* samples included in this study, using the same 84 individuals as in the mitochondrial analyses. To accommodate variable sequencing depth across wild and captive individuals, we used ANGSD to estimate genotype likelihoods rather than hard genotypes. We employed the GATK genotype likelihood model and required high-quality bases and reads by filtering out sites with base quality below 30 or mapping quality below 30, retaining only uniquely mapped reads and excluding reads flagged as problematic. We also performed BAQ recalibration to reduce the influence of misalignments around indels and kept all properly and improperly paired reads to maximize data retention. To ensure reliable allele frequency estimates, we required a minimum representation of half of individuals per site (*-minInd 52*) and applied per-individual depth filters, retaining only those sites with an individual sequencing depth between 3× and 70×. We additionally restricted analyses to biallelic SNPs with a minor allele frequency of at least 5% and identified variable sites using a stringent SNP p-value threshold of 1×10□□. Finally, we computed major and minor alleles, allele frequencies, and genotype likelihoods, exporting the latter in Beagle format for downstream analyses. Then, we used ngsDist (Vieira et al., 2015) to generate a distance matrix based on the genotype likelihoods (*--probs --n_boot_rep 1000 --boot_block_size 500*). We inferred a phylogeny from a bootstrapped distance matrix with fastME (Lefort et al., 2015) and the branch support values were assigned to the final tree using IQ-TREE 2 (see Figure S1).

We next set out to infer the *Pyrrhura* nuclear species tree. To ensure full taxonomic coverage, we selected 33 *Pyrrhura* specimens representing all 24 accepted *Pyrrhura* species and eight recognized subspecies. We included two wild *P. lepida* individuals to account for possible intraspecific complexity because they exhibited divergent mitogenome placements (see Table S2 for details). For each sample, a consensus sequence excluding the sex chromosome was generated with ANGSD v0.94 using the option *-doFasta 3*, applying a minimum mapping quality (--*minMapQ 30*) and base quality (*--minQ 30*) of 30 and requiring a minimum depth of 10 to call a genotype. We additionally removed low-quality or problematic reads, restricted analyses to use only uniquely mapped reads, and required proper read pairing. Consensus sequences were output as compressed FASTA files. Then, we defined non-overlapping genomic windows of 25 kbp. For each of these loci, we extracted the corresponding region from each sample’s consensus genome sequence using samtools and concatenated all sequences for that window into a single multi-FASTA file. Windows were further filtered to retain only high-quality loci by keeping only sequences with less than 10% missing data and loci containing data from at least 95% of the samples. The remaining 1,392 25-kbp alignments were used for inference of the species-tree. For this, we first inferred a maximum-likelihood gene tree for each locus using IQ-TREE 2 (Minh et al., 2020), specifying the GTR+F+G4 substitution model and computing branch support using 1,000 ultrafast bootstrap replicates (*--ufboot*) and 1,000 SH-aLRT tests. Gene trees were filtered to collapse branches with lengths less than 1×10^-5^ substitutions/site and clades with support <1 using the *di2multi* function in the R package *ape* (<u>10.32614/CRAN.package.ape</u>). Finally, we used ASTRAL-III (Zhang et al., 2018) to infer the *Pyrrhura* nuclear species tree, employing the maximum-likelihood gene trees as input. Node support was assessed using 100 replicates of multi-locus bootstrapping to generate quartet support values and posterior probabilities.

## Results

### Dataset characteristics

We generated whole-genome sequencing data for a total of 78 *Pyrrhura* samples. After quality filtering (≥Q30), coverage ranged from 0.68× to 31.5× in the whole data set. Among it, 53 individuals were sequenced to a low-coverage (< 10×) ranging from 0.68× to 9.94× (mean 5.86×). Complete mapping statistics including number of reads, endogenous and clonality can be found in Table S2.

We assembled complete, circular mitogenomes from 75 of the 78 *Pyrrhura* samples and six outgroup samples. *Pyrrhura* mitochondrial genomes range from 16,976 to 17,001 bp, with that of *Pyrrhura pfrimeri* being the smallest. These differences are mainly caused by indels in the control region or other non-coding regions, while the length of protein-coding genes did not exhibit any indels (Table S2). *Pyrrhura* mitochondrial genomes have 22 t-RNA sequences, 2 ribosomal (rRNA) genes, and 13 protein-coding genes. The gene arrangements found in all *Pyrrhura* mitogenomes were uniform and follow the typical avian gene order as previously described for *Pyrrhura* (Urantowka et al., 2016). This arrangement is typical among Arini species (Eberhard & Wright, 2016). The GC content of the complete *Pyrrhura* mitochondrial sequences ranges from 47.2% to 47.7%, relatively high among parrots, which typically have values between 43.2% and 48.9% (Eberhard & Wright, 2016). Mean pairwise identity was calculated to be 96.4% among *Pyrrhura* sequences, and overall 75.3% of sites were identical. The alignment matrix of 17,144 bp included complete mitochondrial genomes of *Pyrrhura* samples, as well as outgroup sequences.

### Phylogenetic inference

#### Nuclear genome

All *Pyrrhura* species grouped in two monophyletic and highly supported clades, similar to Smith et al. (2023), here referred to as nuclear clade 1 (Nr-1) and nuclear clade 2 (Nr-2) (Figure 1). Nr-1 is composed by the classic *picta-leucotis* complex of species, including *Pyrrhura cruentata* in a basal position to it.

**Figure 1.**
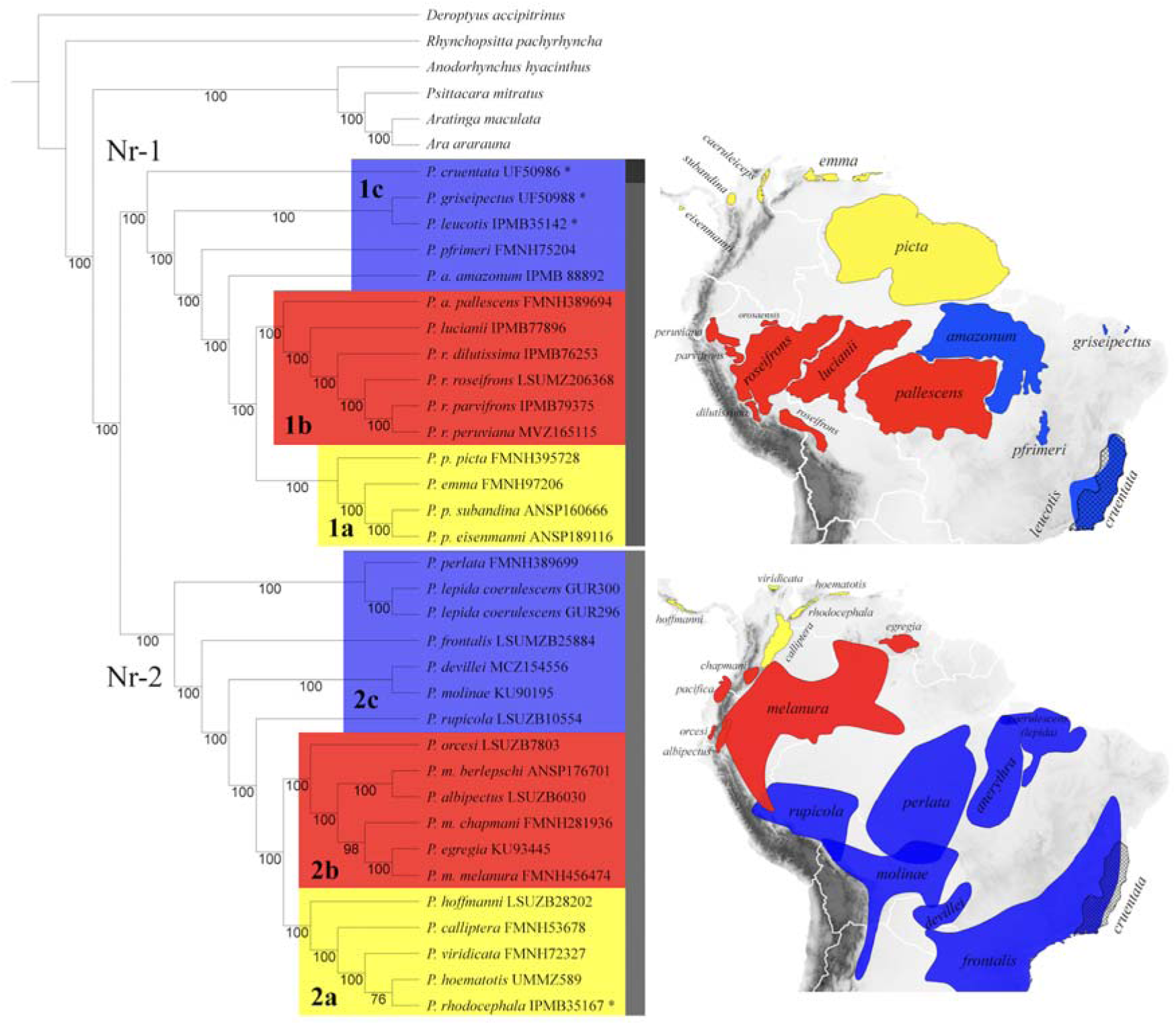
Nuclear-genome phylogeny, based on 1392 alignments, inferred by ASTRAL-III, and distribution of selected *Pyrrhura* species and subspecies. Grey bars indicate the classic *Pyrrhura* groups formed using mitochondrial DNA: *P. cruentata,* the *picta-leucotis* complex, and the rest of the species (here, *rupicula-melanura*). The ASTRAL tree reveals two clades: Nuclear-Clade 1 (Nr-1; picta-leucotis complex) and Nuclear-Clade 2 (Nr-2; the rest of *Pyrrhura* species). Both nuclear clades can be subdivided into three main groups and show similar geographical patterns: A monophyletic clade of mainly restricted-range species associated to the Northern Andes (in yellow); a sister monophyletic clade of restricted-ranged species associated to the Andes and widespread species associated to the Amazon (in red); and a paraphyletic clade of early divergent species (in blue). Node labels indicate percent support from bootstrap replicates. Asterices (*) next to the species label highlight that the taxon is represented by a captive sample.

Within Nr-1, we observe three clear groups: a monophyletic clade (here referred to as Nr-1a) containing *P. emma*, *P. picta*, *P. p. subandina*, and *P. p. caeruleiceps*; a sister monophyletic clade (Nr-1b) containing *P. roseifrons*, *P. r. peruviana*, *P. r. parvifrons*, *P. r. dilutissima*, *P. lucianii*, and *P. a. pallescens*; and a paraphyletic group (Nr-1c) containing the remaining species *P. amazonum*, *P. pfrimeri*, *P. griseipectus*, and *P. leucotis* that diverged earlier from the other two clades. Within Nr-2, we observed similar diversification patterns as three groups were formed: the first monophyletic clade (Nr-2a) contains *P. rhodocephala*, *P. hoematotis*, *P. calliptera*, *P. hoffmanni*, and *P. viridicata*; the second monophyletic clade (Nr-2b, sister to Nr-2a) contains *P. egregia*, *P. melanura* and related taxa, *P. albipectus*, and *P. orcesi*; and the third paraphyletic group (Nr-2c) contains the remaining species that diverged earlier from Nr-2a and Nr-2b. Within Nr-2c, *P. perlata* and *P. lepida* form a clade, as do *P. molinae* and *P. devillei*. For *P. melanura*, regarded as a species complex, we observed deep divergences among taxa, especially for *P. melanura berlepschi*, which seems to be closer to *P. albipectus* than to the other *P. melanura* samples.

#### Mitochondrial genome and mito-nuclear discordances

The maximum-likelihood (Figure 2) and Bayesian (Figure 3, Figure S2) phylogenies based on entire mitogenome sequences showed high support for most species-level clades with bootstrap values over 85/100 and posterior probabilities (PP) over 0.97. However, we found low support (PP = 0.56) for the relationship between *P. roseifrons* and its sister clade, as well as for the relationship between *P. egregia* and its sister clade (PP = 0.66). We did not find topological disagreements between the methods, and both maintained the three classic and highly supported groups of *Pyrrhura* species: (a) *Pyrrhura cruentata*, (b) the *picta-leucotis* complex, and (c) the rest of the species (Ribas et al. 2006).

**Figure 2.**
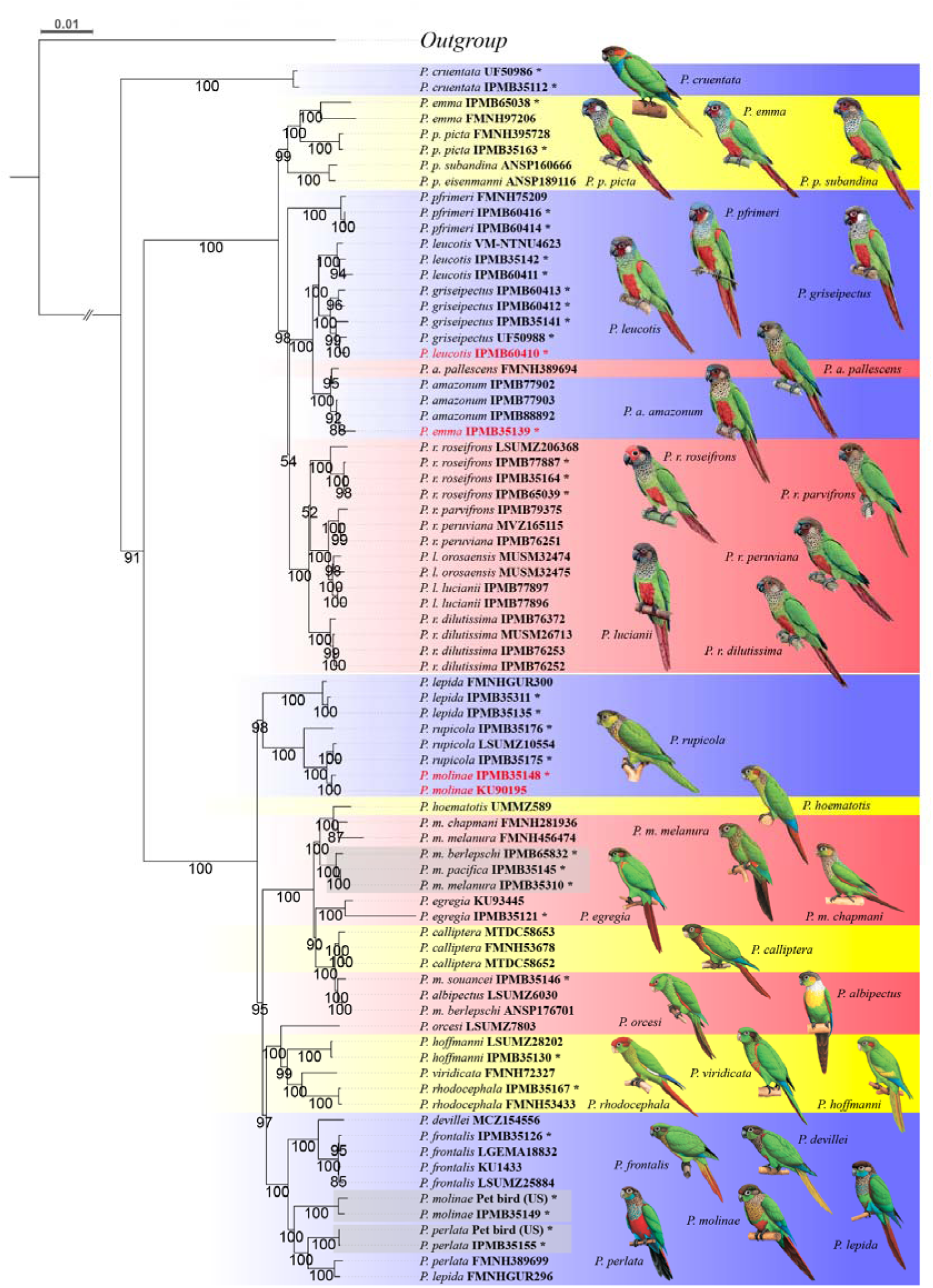
Maximum-likelihood phylogeny of ∼16,990 bp corresponding to complete mitochondrial genomes. The mitogenomes of species within nuclear clade 1 (Nr-1), correspond to the *picta-leucotis* complex and *P. cruentata* clade. While the mitogenomes of the species within Nr-2 form a separated clade. Captive individuals are noted with an asterisk (*), unexpected placements are labeled in red, and grey backgrounds highlight three captive clades. Clades are colored following the three subclades formed on the nuclear tree (blue = early divergent species, red and yellow = monophyletic clades). *Pyrrhura* color plates reproduced with the permission of Lynx Editions. Ultrafast bootstrap support values are indicated at the nodes.

**Figure 3.**
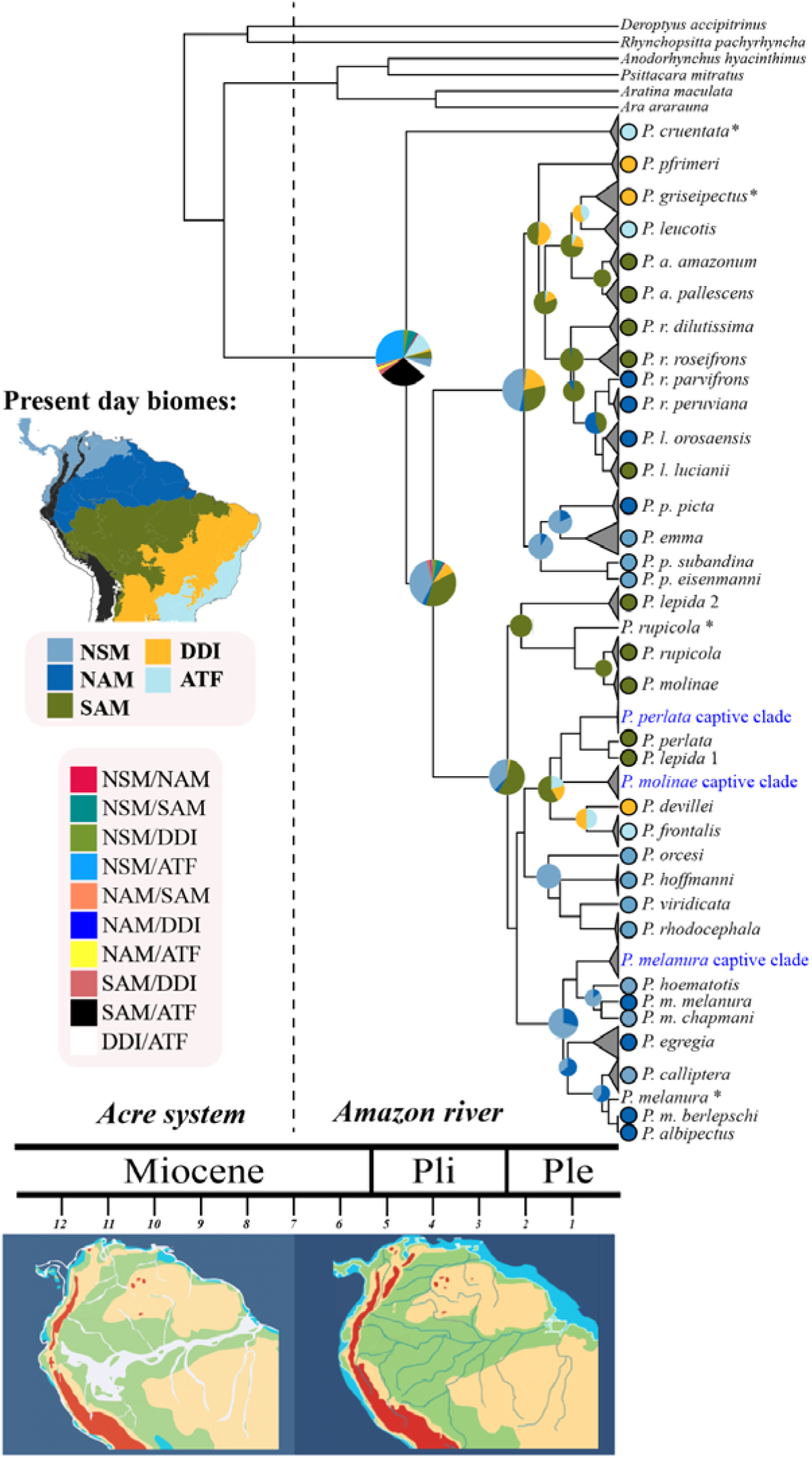
Ancestral range reconstruction and time-calibrated phylogeny of the genus *Pyrrhura*. The five present-day biomes based on Morrone et al. (2022) are shown, the Andes mountains are shown in black. Colored circles next to the species labels represent the assigned biogeographical regions considered in the analysis. All collapsed clades, with the exception of the “captive clades” (shown in blue), contain at least one wild specimen, except for *P. cruentata* and *P. griseipectus.* Captive individuals (marked with an asterisk) and captive clades (in blue) were excluded from the biogeographic analysis. Pie charts on the nodes represent the speciation area inferred by BioGeoBEARS (DIVALIKE+J). Graphic illustrations of the geographical evolution of the Neotropical region are based on Hoorn et al. (2010), and correspond to the time period they are on.

Despite this, we observed several unexpected placements. First, we identified a captive *P. leucotis* individual (IPMB60410) clustering within the *P. griseipectus* mitogenomic clade, and a captive *P. emma* individual (IPMB65038) clustering within the *P. amazonum* clade. However, analysis of nuclear genomic data (Figure S1) showed that both specimens cluster with individuals of their respective nominal species. This mito–nuclear discordance indicates that these individuals are of hybrid origin. On the other hand, we unexpectedly observed two *P. molinae* (IPMB35148, KU90195), a captive and a wild specimen, placed together within the *P. rupicola* clade (Figure 2A). In addition, we identified three distinct clades composed exclusively of captive specimens, each corresponding to a single nominal species (*P. melanura*, *P. molinae*, and *P. perlata*). These captive clades show substantial genetic divergence from the clades containing wild individuals of the same species (Table S3). However, nuclear genomic analyses indicate a closer relationship between captive and wild individuals within each nominal species (Figure S1).

Interestingly, even after excluding captive individuals, we observed two distinct placements for *P. lepida*; one sample grouped with *P. perlata* while three samples, including a wild specimen, formed a clade sister to *P. rupicula* (Figure 2). The latter placement has not been previously described. However, samples carrying these two mitochondrial lineages of *P. lepida* clustered in the nuclear tree as clade sister to *P. perlata* (Figure 1). Similarly, even considering only wild individuals, we found that *P. melanura* is paraphyletic in the mitogenome based tree (Figure 2), and in the nuclear tree (Figure 1).

The mitogenome and nuclear phylogenies recovered the same two major clades, with the exception of *P. cruentata*. In the mitochondrial tree, *P. cruentata* appears as sister to all other *Pyrrhura* (Figure 2), whereas in the nuclear phylogeny it occupies a basal position within Nr-1 (Figure 1). This discordance indicates the presence of two major nuclear lineages (Nr-1 and Nr-2) but three mitochondrial lineages. Setting *P. cruentata* aside, we observed that the rest of Nr-1c mitogenomes were not found in a basal position as in the nuclear tree but rather closer to species corresponding to Nr-1b (Figure 2). Controversily, the mitogenomes of the monophyletic clades Nr-1a and Nr-1b also form well-supported monophyletic groups, broadly corresponding with the nuclear phylogeny. An exception is *P. amazonum pallescens*, whose mitogenome clusters with nominal *P. amazonum* individuals (Nr-1c) rather than with Nr-1b. In addition, we also observed that the mitogenome of *P. lucianii* was placed within the *P. roseifrons* clade rather than sister to them, having a closer relationship with the geographically distant subspecies *P. r. parvifrons* and *P. r. peruviana*.

Within Nr-2, we observed that species belonging to the basal Nr-2c lineage segregate into two distinct monophyletic clades (Figure 2). One clade retains a basal position relative to Nr-2a and Nr-2b, while the other is closely related to *P. hoffmanni* and its allies (Nr-2a). In addition, the mitogenomes of most Nr-2a and Nr-2b taxa also form respective monophyletic clades. However, we also observed disagreements. For example, *P. calliptera* and *P. hoematotis*, which fall within Nr-2a in the nuclear phylogeny, instead group with Nr-2b species in the mitochondrial tree. Conversely, the mitogenome of *P. orcesi* appears as sister to *P. hoffmanni*, *P. viridicata*, and *P. rhodocephala* (Nr-2a), rather than to the remaining Nr-2b taxa, as indicated by the nuclear data. Another important discordance is the existence of two wild *P. lepida* mitochondrial lineage where one is placed congruently with *P. perlata*, but the remaining one is placed closer to *P. rupicola*, but geographically distant.

#### Biogeography, ancestral ranges and divergence times

Based on our time-calibrated mitogenome phylogeny (Figure 3) and ancestral range reconstructions using BioGeoBEARS, we found that the most recent common ancestor of *Pyrrhura* diverged from other *Arini* species around 8.68 Mya, during the late Miocene (Figure 3). The mitochondrial lineage of *P. cruentata* split from the rest of *Pyrrhura* approximately 4.66 Mya, followed shortly after by the divergence of the remaining *Pyrrhura* into two major mitochondrial lineages at around 4.08 Mya. We observed that the monophyletic clades Nr-1a and Nr-2a consist primarily of species with relatively small geographic ranges distributed across northern South America and Mesoamerica (NSM). Except for *P. picta*, all taxa in these clades occur in or near the Northern Andes and show strong associations with particular elevation bands, resulting in largely non-overlapping distributions. Our BEAST analysis indicates that the diversification of the mitogenomes lineages corresponding to Nr-1a and Nr-2a occurred relatively recently, around 1.73 Mya [95% HPD, 0.9-2.8] and 1.3 Mya [95% HPD, 0.6-2.1], respectively, both in northern South America. In contrast, taxa assigned to Nr-1b and Nr-2b are primarily associated with the western and central Amazon. Although some taxa, such as *P. melanura*, occupy wide ranges across the Amazon Basin, most exhibit smaller and more localized distributions, often along the foothills of the Central Andes. Our BEAST analysis estimates that the mitogenome lineages of all Nr-1b taxa excluding *P. a. pallescens* originated approximately 1.1 Mya [95% HPD, 0.5-1.7], most likely in the southern Amazon (SAM). Similarly, the mitogenomes of most of Nr-2b originated around 1.2 Mya [95% HPD, 0.6-2], likely in the northern Amazon (NAM).

## Discussion

### Evolutionary relationships within *Pyrrhura*

Our study presents the most comprehensive phylogenetic framework for *Pyrrhura* to date, integrating nuclear genomic data with densely sampled mitogenomes across all recognized species and several other taxa. The resulting nuclear phylogeny is strongly supported at all nodes, and although it is broadly concordant with the UCE-based phylogeny of Smith et al. (2023), we found important disagreements (Figure S3). For instance, we found high support for *P. amazonum* forming a distinct lineage sister to the Nr-1a and Nr-1b clades, instead of placed with low support within Nr-1b. Similarly, our dataset provides strong support for the sister relationship between *P. leucotis* and *P. griseipectus*, a relationship poorly resolved in Smith et al. (2023). We also observed a high support for the basal position of *P. orcesi* within Nr-2b, rather than a low-supported position sister to both Nr-2b and Nr-2a, and high support for the position of *P. hoffmanni* and *P. calliptera* which appear in opposite placements and with low support in the UCE phylogeny. Our extensive sampling size, that includes many intraspecific taxa, could explain the differences, particularly the inclusion of *P. pfrimeri*, the only species absent in Smith et al. (2023), could have minimized the risk of long branch attraction between the early divergent species. In addition, the chosen genomic approach could have played a role as we used whole genome sequence data instead of UCEs, which could limit the study’s power to resolve evolutionary relationships in the face of rapid radiation and hybridization.

We also observed *P. devillei* to be sister to *P. molinae*, whereas Smith et al. (2023) recovered *P. devillei* as sister to *P. frontalis*. However, despite these differences, both topologies are well supported. Although, little is known about *P. devillei* at the molecular level, it has been considered closer to *P. frontalis* (specifically *P. frontalis chiripepe*) (Collar, Boesman, et al., 2020), aligning with the findings of Smith et al. (2023). However, it is necessary to consider that *P. devillei* is distributed between the ranges of *P. frontalis* and *P. molinae*, with potential overlaps (Figure 1) and recorded hybridization in northern Paraguay (Collar & Boesman, 2020), suggesting that topological discordances could be attributed to the sampling localities. On their study, *P. devillei* (AMNH 785882) was originally collected in Puerto Guaraní, Paraguay, and *P. molinae* (DOT 2706) came from Nor Yungas (Bolivia), while *P. frontalis* (LSU B-14206) was represented by a captive individual. In contrast, our nuclear phylogeny included a *P. devillei* (MCZ 154556) specimen from Miranda, Brazil, a *P. molinae* (KU 90195) from Picada Chovoreca, Paraguay, and a *P. frontalis* (LSU B-25884) from Caazapá, Paraguay. This means that our phylogeny includes a *P. devillei* sample from a place where there are no records of inter-specific gene flow. Moreover, while Smith et al. (2023) compared his *P. devillei* data with a geographically distant *P. molinae*, we compare ours with nearby populations of both *P. frontalis* and *P. molinae*. Therefore, our results are not only the best hypothesis for *P. devillei* position within *Pyrrhura*, but also represent more closely these interaction zones and emphasizes the importance of accounting for intraspecific structure in phylogenetic analyses.

### Biogeography and temporal diversification of *Pyrrhura*

Ancestral range inference supports a southern origin for the genus *Pyrrhura*, with the most recent common ancestor diverging from other Arini around 8.7 Mya during the late Miocene (Figure 3). This timing coincides with the existence of the Acre system, a vast wetland complex that divided the northern and southern portions of the Amazon Basin between ∼10 and 7 Mya (Hoorn et al., 2010). Comparable patterns of early Amazonian diversification driven by the Acre and earlier Pebas systems have been documented in other groups such as butterflies (Condamine et al., 2012), and frogs (Ortiz et al., 2023). Given that northern Amazonian lineages of *Pyrrhura* appear only much later in the phylogeny ca. 2 Mya, our results support a scenario in which the genus originated south of the Amazon River before subsequently dispersing northward. Moreover, the fact that all early-divergent species from both major nuclear clades (Nr-1c and Nr-2c) are geographically restricted to regions south of the Amazon River, coinciding with the hypothetized ancestral range of *Pyrrhura*, provides further support for a southern Amazonian or transitional Atlantic Forest origin of the genus. Moreover, as discussed by Ribas et al. (2006), the Atlantic Forest is the only biome where the three major mitochondrial lineages occur, further supporting this scenario.

Smith et al. (2023) dated the original split between the two major nuclear lineages of *Pyrrhura* to approximately 5.1 Mya [2.2–9.9 Mya]. Although our nuclear results also support two primary lineages (Nr-1 and Nr-2), the mitogenome phylogeny confirms the existence of three mitochondrial lineages: one formed exclusively by *P. cruentata*, and two others corresponding to the nuclear Nr-1 and Nr-2 clades. We estimate that *P. cruentata*’s mitogenome diverged from all other *Pyrrhura* around 4.66 Mya, followed by the split between the remaining mitochondrial lineages at approximately 4.08 Mya. Both divergence times fall well within the confidence interval of the nuclear estimate (Smith et al., 2023), yet they raise the question of why the mitochondrial genome retains three lineages. One possibility is incomplete lineage sorting, where *P. cruentata* retained an ancestral mitochondrial haplotype that did not fully sort before the early diversification of *Pyrrhura*. Alternatively, the mitochondrial lineage found in the rest of Nr-1c species may be reflecting historical introgression among geographically adjacent taxa, making them closer to Nr-1b in the mitogenome tree, while *P. cruentata,* who was isolated in the Atlantic Forest, retained the ancestral mitochondrial lineage of the Nr-1 nuclear clade. This latter scenario is supported by the fact that the Dry Diagonal has been considered a major biogeographic barrier in other Neotropical groups (Bocalini et al., 2021; Reyes et al., 2023).

The diversification of *Pyrrhura* accelerated between 1 and 3 Mya, coinciding with the emergence of two monophyletic sister subclades (“a” and “b”) within both Nr-1 and Nr-2. The diversification of Andean-associated subclades Nr-1a and Nr-2a occurred during the Pleistocene, at approximately 1.7 and 1.3 Mya, respectively. These dates coincide with periods of significant ecological opportunity after the colonization and diversification of high-altitude, species-rich flora between 2 and 4 Mya (Hughes & Eastwood, 2006) as a consequence of the the final phases of Northern Andean uplift between 8 and 5 Mya (Pérez-Escobar et al., 2022). The absence of range overlaps among species of this group, combined with their strong elevational associations, and the lack of obvious morphological or ecological innovations suggests that diversification may have proceeded through an “island-like” process in which isolated montane habitats promoted allopatric speciation. This pattern is similar to what was found in those Andean plants (Hughes & Eastwood, 2006).

In contrast, our nuclear phylogeny suggests that the diversification of the Amazonian subclades Nr-1b and Nr-2b followed a speciation-by-dispersal scenario, as younger species are geographically farther from the estimated ancestral range of *Pyrrhura*. In addition, the association of *Pyrrhura* with the Andes may reflect a response to Pleistocene temperature fluctuations, as birds are known to shift their altitudinal ranges to maintain optimal thermal conditions (Freeman et al., 2018). Such shifts could have led to population isolation, similar to the patterns observed in the Northern Andes. Additionally, the early-diverging species (Nr-1c and Nr-2c) are primarily distributed across the Amazon Basin, with their ranges constrained to the north by the Amazon River. Their parapatric distributions and close association with river systems suggest that their diversification may have been influenced by habitat fragmentation during the Pleistocene, in line with the forest refugium hypothesis (Arndt & Wink, 2017).

### Role and limitations of mitogenomes in *Pyrrhura* phylogeny

Although nuclear genomic data provide the most reliable framework for resolving deep phylogenetic relationships (Tamashiro et al., 2019), the fast-evolving mitochondrial genomes (Kumar, 1996) remain valuable for detecting recent divergence, cryptic diversity, and potential introgression and hybridizations, particularly in groups with such a complex evolutionary histories. In addition, current genomic studies are not yet as extensive as earlier mitochondrial research (Arndt & Wink, 2017; Ribas et al., 2006), which have been the base for several systematic revisions within the genus (Stotz, 2007). Therefore incorporating mitogenome evidence allows us to make cross-study comparisons. Moreover, as mitochondrial data is relatively easier to generate, it allows us to wider the sampling size, which is particularly important for mitochondrial-based phylogenetic inference as shown in tree squirrels (de Abreu-Jr et al., 2020), another rapidly evolving Neotropical vertebrate lineage.

Despite this, it is important to consider currently that up to 5% of published bird mitochondrial genomes are problematic due to misidentification, chimeras of multiple species, nuclear-mitochondrial insertions, and sequencing errors (Sangster & Luksenburg, 2021). Therefore, to avoid incorrect phylogenetic hypotheses, careful assembly, annotation, and interpretation are essential. Before the present work, only a single *Pyrrhura* mitochondrial genome was published in GenBank (NCBI), derived from a captive hybrid specimen (Urantowka et al., 2016). However, Our study incorporates complete mitochondrial genomes for 75 *Pyrrhura* specimens, representing the broadest mitogenomic sampling to date for the genus, and increasing our confidence in the results.

The mitogenome-based phylogeny recovered the three well-supported mitochondrial lineages originally identified by Ribas et al. (2006). However, in contrast to their results, we found *P. cruentata* to be sister to all *Pyrrhura* lineages rather than sister only to the *picta–leucotis* complex. Within the *picta–leucotis* group, Ribas et al. (2006) proposed four major clades, whereas Arndt & Wink (2017) suggested up to six. Our results strongly support the former arrangement (Figure 2) and provide improved resolution of the relationships among these clades. Similarly, mitogenomes of Nr-2 species formed four major clades largely consistent with Ribas et al. (2006), and our dataset yielded greater statistical support for their interrelationships. For example, the placement of *P. pfrimeri*, previously difficult to resolve, was strongly supported in our analysis as sister to the clade containing *P. roseifrons*, *P. leucotis*, and related taxa (PP = 1.0, 98/100 bootstraps).

### Mito–nuclear discordances and its underlying causes

Discordances between mitochondrial and nuclear phylogenies are common in birds (Campillo et al., 2019; Noll et al., 2022; Pavlova et al., 2013) and may arise from hybridization, which is frequently observed even under apparent selection against it (Blom et al., 2024). Particularly in parrots there are documented examples of hybridization even among non-congeneric individuals (Hingston, 2022). Moreover, given that mitogenomes do not undergo recombination, they are more susceptible to incomplete lineage sorting (ILS), historical or ongoing hybridization, introgression, or even selection on female traits (Morales et al., 2015), especially under rapid radiation scenarios as in *Pyrrhura* where more than 30 lineages appeared over the course of 5 million years.

The most prominent mito-nuclear discordance involves *P. cruentata*, whose mitogenome forms a deeply divergent lineage sister to all other *Pyrrhura*, while its nuclear genome places it basal within Nr-1. As discussed above, this conflict may reflect retention of an ancestral mitochondrial haplotype through incomplete lineage sorting (ILS) or long-term isolation of Atlantic Forest populations that prevented the mitochondrial introgression observed in other Nr-1 species. Because *P. cruentata* is endemic to the Atlantic Forest, positive selection could also have contributed to the fixation of its mitochondrial lineage following divergence, a process documented in other avian groups such as penguins (Noll et al., 2022). Additionally, the pronounced morphological differences of *P. cruentata* relative to other Nr-1 taxa may reinforce pre-zygotic isolation, limiting hybridization with *P. frontalis* and *P. leucotis* even where their distributions overlap. Such barriers, combined with long-term ecological isolation, could explain why no hybridization has been reported in the wild among these species.

Similarly, the early divergent species of Nr-2 (Nr-2c) form two monophyletic clades in the mitochondrial phylogeny. One of these lineages, comprising *P. rupicola* and allies, occupies a basal position relative to all other Nr-2 species, with an estimated divergence of ∼3.05 Mya, consistent with the diversification times of Nr-2c reported by Smith et al. (2023). This suggests that it represents the ancestral mitochondrial lineage of the clade. In contrast, the second mitochondrial lineage of Nr-2c (P. lepida and related taxa, see figure 2) clusters more closely with the monophyletic Nr-2a group, despite the expectation that geographic proximity should favor affinity with Nr-2b. These contrasting patterns highlight two major classes of mito–nuclear discordance. First, mitochondrial similarity between geographically distant taxa may reflect retention of ancestral mitochondrial haplotypes, lineage-specific fixation through drift or selection, or other deep-time evolutionary processes. Second, mitochondrial similarity among geographically close species, despite deeper nuclear divergence, can be more readily explained by historical gene flow or mitochondrial introgression. For example *P. amazonum pallescens* shares mitochondrial affinity with nominal *P. amazonum* rather than with Nr-1b, consistent with recent or past gene flow between neighboring populations (Figure S1).

### Detecting overlooked genetic diversity by incorporating captive specimens

Captive individuals may occasionally be misidentified or descend from undocumented hybridization events. In this study, we identified two such hybrids: a *P. leucotis* x *P. griseipectus* (IPMB60410), and a *P. emma* x *P. amazonum* sample (IPMB65038). These specimens originated from Loro Parque (Tenerife, Spain), where staff take considerable care to prevent interspecific hybridization and avoid working with morphotypes commonly produced in the pet trade. However, morphological identification of taxa such as *Pyrrhura emma* can be challenging, especially for non-specialists, and it cannot be excluded that intentional or unintentional hybridization occurred prior to their arrival at Loro Parque. According to staff reports, some birds may have originated from South American breeding centers three to six generations earlier, where hybridization could have taken place (Rafael Zamora, personal communication).

This reinforces the need to interpret captive-derived data with caution. Nevertheless, when evaluated alongside nuclear evidence and specimens from the wild, captive samples can be highly informative for detecting cryptic genetic diversity. In our study, the inclusion of captive and museum material allowed us to obtain mitochondrial genomes for taxa that are otherwise difficult to sample, and in several cases revealed unexpected and previously overlooked genetic structure. For example, we identified a highly divergent captive *P. rupicola* specimen (IPMB 35176) whose mitogenome shows levels of divergence comparable to those observed between recognized species (Figure 2). Although originally identified as *P. r. sandiae*, a subspecies currently considered undiagnosable (Collar, A., et al., 2020) but recognized (Collar, A., et al., 2020), the documented morphological variation (Bond & Meyer de Shauensee, 1944) and our mitogenomic results support complex evolutionary history. Nuclear data confirm its assignment to *P. rupicola*, indicating the presence of deep intraspecific structure rather than species-level divergence. Similarly, we detected substantial mitochondrial differentiation within *P. emma* and *P. egregia* (Table S3), consistent with additional taxonomic complexity, whether already recognized (e.g. *P. egregia obscura*) or still unrecognized (e.g. *P. emma auricularis;* Zimmer & Phelps 1949). These findings underscore the value of broad geographic sampling, across both wild and captive sources, for accurately characterizing evolutionary diversity within *Pyrrhura*.

We also detected three divergent captive lineages in the mitogenome phylogeny (Figure 2), each consisting exclusively of captive-bred individuals of *P. melanura*, *P. molinae*, and *P. perlata*. The mitochondrial captive clade of *P. melanura* (*n* = 3) is close to the clade that includes wild *P. m. melanura*, *P. m. chapmani*, and *P. hoematotis*. However, two of these captive individuals do not group with *P. melanura* in the nuclear tree and rather cluster with *P. orcesi* (Figure S1). This discordance is consistent with the previously discussed taxonomic and evolutionary complexity of the *P. melanura* group. In contrast, both the *P. molinae* (n = 2) and the *P. perlata* (n = 2) captive clades were found closely related to the wild *P. perlata* and *P. lepida* (Figure 2). These captive individuals cluster with all other captive and wild specimens of the corresponding species (Figure S1). Notably, in both cases the two individuals forming these captive lineages originated from different countries (Spain and the United States), suggesting that their shared mitochondrial lineages reflect genuine biological variation rather than localized breeding-facility effects. These findings indicate that captive birds can harbor mitogenomic diversity consistent with complex intraspecific structure within *P. molinae (Collar & Boesman, 2020)*, and the *Pyrrhura lepida-perlata* complex (Collar & Boesman, 2020; Somenzari & Silveira, 2015).

### Implications for *Pyrrhura* systematics

We have discussed that some of the topological differences between phylogenetic inferences could be attributed to the sampling locality chosen to represent the species, particularly when great intra-specific variation is possible. For instance, the specimen of *P. amazonum* (LACM 42239) used by Smith et al. (2023) was originally sourced in Imperatriz (Tocantins River) while the samples we utilized were sourced from Altamira (Xingu River) and Santana do Araguaia (Araguaia River). These localities are far from the species’ type locality (Obidos, west of Tapajos River), and fall into the distribution area of two proposed related taxa “*P. a. microtera*” (Todd, 1947) and “*P. a. araguaiensis*”. This means that, although these taxa are not recognized, the potential nominal *P. amazonum* might have not been included in this or previous phylogenies (Arndt & Wink, 2017; Ribas et al., 2006) and future studies should try to include specimens to investigate whether there are grounds to differentiate these taxa.

A related systematic question is whether *P. amazonum pallescens,* considered a subspecies of *P. amazonum* by the IOC, should be considered as a full species, including *melanoides* (=*lucida*) as its subspecies, or whether both taxa should remain as subspecies of *P. amazonum*. Previous molecular work could not separate *P. a. amazonum* from *P. a. pallescens* (Arndt & Wink, 2017; Ribas et al., 2006). Particularly, Arndt & Wink (2017) reported clear morphological, ecological, vocal, and ethological differences but could not differentiate them based on Cyt-b gene alone. Our mitogenome analysis discriminates between our wild *P. a. pallescens* specimen from the rest of *P. a. amazonum*. Moreover, our nuclear analysis revealed significant genomic divergences between these two taxa for the first time, further supporting their split.

Historically, *P. picta* encompassed a great variety of taxa that were split in light of molecular evidence (Ribas et al., 2006). However, despite clear geographical differentiation, some taxa such as *P. p. eisenmanni*, *P. p. caeruleiceps,* and *P. p. subandina* are still regarded as subspecies of *P. picta* (Gill et al., 2025) due to their weak morphological differentiation (Collar, del Hoyo, et al., 2020). *P. emma*, on the other hand, is accepted by the IOC but not by SACC. Our nuclear analysis corroborates that there is sufficient genomic differentiation among these taxa (Smith et al., 2023), with *P. picta* and *P. emma* placed basal to the rest of Nr-1a species. Moreover, our mitogenome analysis corroborates the differentiation of *P. emma* and *P. picta* from the rest within the group, aligning with their geographical distributions. Although the divergence within this group is fairly recent (around 1.73 Mya), it is older than the split between *P. leucotis* and *P. griseipectus*, and even older than the split of *P. leucotis*/*P. griseipectus* from *P. amazonum* (nearest neighbour to the clade). In summary, current taxonomy distinguishes two species *P. emma* and *P. picta.* However, according to our analysis the latter is clearly paraphyletic, thus a taxonomic re-organization of this complex is necessary. This would either treat all forms as conspecific under *P. picta* (Müller, 1776), which has nomenclatural priority and would include *P. p. emma*, or grant species status to all those taxa in geographical isolation, distinguishing five species in total.

We also address the status of *P. roseifrons*, currently regarded as a full species by the IOC, which includes the following taxa: *P. r. peruviana*, *P. r. parvifrons*, and *P. r. dilutissima,* being *P. r. dilutissima* often considered to be closer related to *P. r. peruviana* (del Hoyo et al., 2022). Our analysis shows that nuclear and mitochondrial trees agree on the same topology: (((*P. r. peruviana*, *P. r. parvifrons*), *P. r. roseifrons*), *P. r. dilutissima*) (Figure S4). This arrangement and our ancestral geographic range analysis suggest a directional isolation by distance along the Andes from south to north, starting a transition from Amazonian *Pyrrhura* to montane *Pyrrhura*. Although our data set is limited, we consider that there is enough evidence to at least separate *P. r. peruviana/P. r. parvifrons* from *P. roseifrons*. Moreover, in the nuclear tree *P. r. dilutissima* was placed sister to all the taxa in the group and not sister to only *P. r. peruviana* as expected by morphological similarities (del Hoyo et al., 2022). The genetic differentiation and geographical isolation from all other *P. roseifrons* taxa further support the status of *P. r. dilutissima* as a full species.

Regarding the specimens from the Orosa River, here referred as *P. l. orosaensis*, which were proposed to be a new subspecies of the otherwise monotypic *P. lucianii* (Arndt & Wink, 2017). Not only there is a big geographical disconnection between the ranges of *P. l. orosaensis* and the nominal *P. leucotis*, but our results, including wild individuals from Natural History Museum of San Marcos (MUSM, Peru), shows that the genetic distances between *P. l. orosaensis* and *P. lucianii* in the mitogenome tree are comparable to the distances separating accepted subspecies (e.g. *P. r. peruviana* and *P. r. parvifrons*) (Table S3). This supports the existence of *P. l. orosaensis*, which would need further morphological analysis.

Within nuclear clade Nr-2, few comprehensive molecular studies have addressed the longstanding systematic challenges, particularly challenging within the *P. melanura* complex and the *P. perlata–lepida* group. For starters, *Pyrrhura melanura* has long been suspected to represent a species complex; for example, *P. m. pacifica* was proposed to have greater affinity with *P. orcesi* than with other *P. melanura* taxa (Ridgely & Robbins, 1988). However, due to the historical lack of broad molecular sampling, proposed splits from the nominal form have not been widely accepted (Penhallurick, 2012), and most phylogenetic studies have relied on a single specimen to represent the entire species. *Pyrrhura melanura* is one of the few *Pyrrhura* taxa occurring north of the Amazon River with a wide geographic distribution, and overlooking its internal diversity can lead to misleading phylogenetic interpretations. In our analyses, *P. melanura* is not monophyletic in either the nuclear or mitochondrial phylogeny. Mitogenomes from wild individuals fall into two distinct lineages, consistent with Ribas et al. (2006). Additionally, the captive *P. melanura* clade diverged from the closest wild lineage approximately 0.93 Mya, and another captive individual (IPMB 35143) was recovered as deeply divergent from the clade formed by wild *P. melanura* and *P. albipectus*. These patterns strongly indicate that *P. melanura* harbors substantial cryptic structure. A comprehensive genomic study with dense sampling across the species’ range will be necessary to resolve its taxonomy and assess whether multiple species-level lineages are present.

The *P. perlata–lepida* group presents persistent taxonomic challenges. The IOC currently recognizes *Pyrrhura perlata* and *P. lepida* as full species, with *P. lepida* comprising three subspecies: *P. l. lepida*, *P. l. anerythra*, and *P. l. coerulescens*. Based on a morphological analysis of 174 specimens, Somenzari & Silveira (2015) proposed reducing the complex to three diagnosable taxa: *P. perlata*, *P. anerythra*, and *P. coerulescens*, the latter encompassing what is traditionally treated as *P. lepida*. Our nuclear dataset, which includes two wild *P. lepida* individuals (GUR 296 and GUR 300), one wild *P. perlata* (FMNH 389699), and multiple captive representatives, recovered *P. perlata* and *P. lepida* as two reciprocally monophyletic sister clades (Figure 1 and S1). In contrast, the mitochondrial phylogeny shows strong discordance: GUR 296 clusters with *P. perlata*, as expected, whereas GUR 300 and two captive *P. lepida* specimens form a clade sister to *P. rupicola*. The later placement is unexpected as previous mitochondrial-based studies (Arndt & Wink, 2017; Ribas et al., 2006) generally recovered *P. lepida* as closely related to *P. perlata* and *P. molinae*, with *P. rupicola* forming its own lineage.

These inconsistencies are consistent with unresolved taxonomic boundaries within the *P. lepida–perlata* complex (Brito et al., 2016; Somenzari & Silveira, 2015). Therefore, part of this discrepancy likely reflects mismatched geographic sampling. Following Somenzari & Silveira (2015), which restrict *P. lepida* sensu stricto to *P. coerulescens*, the mitochondrial sample used by Ribas et al. (2006) originates from within the range of *P. anerythra*, whereas our wild samples are from the distribution of *P. coerulescens*. Without sampling *P. anerythra*, we cannot fully evaluate the three-way division proposed by these authors. Nonetheless, our results underscore the importance of multiple representatives per taxon and reveal substantial mito–nuclear discordance indicative of historical introgression or incomplete lineage sorting. A comprehensive genomic reassessment of this complex is needed, and future phylogenetic studies should explicitly incorporate the deep divergence and geographic structuring documented here.

### Implications for the conservation of the genus and future directions

The systematics of *Pyrrhura* parrots are inherently complex due to subtle morphological differentiation and the scarcity of comprehensive taxonomic studies. Although stable nomenclature is essential for governmental conservation agencies, taxonomic revisions in *Pyrrhura* have not kept pace with conservation needs. In particular, several broadly distributed polytypic species such as *P. roseifrons*, *P. melanura*, and *P. molinae*, contain geographically restricted subspecies, many of which occur along the Andes. These taxa are currently assessed as single units and therefore classified as not globally threatened (CITES Appendix II), whereas most *Pyrrhura* species with small distributions are categorized as threatened to varying degrees. Systematic re-evaluation of these polytypic species could reveal that certain isolated populations face substantially higher extinction risks than their current taxonomic status suggests. Because parrots are charismatic, recognizing genetically distinct lineages within *Pyrrhura* can also guide the designation of new conservation areas tailored to preserve evolutionary diversity.

The Andean region, in particular, merits focused investigation. Several ranges that underwent final uplift during the Pleistocene, such as the Subandes and the Sierras Pampeanas (Pérez-Escobar et al., 2022) harbor lineages whose evolutionary histories remain poorly understood. The Subandes may represent a hybridization zone between *P. rupicola* and *P. melanura* (Robbins et al., 2013), and previous work suggests the possible presence of an undescribed *P. roseifrons* lineage in the same region (Arndt & Wink, 2017). Notably, the politypic *P. molinae* spans both the Subandes and Sierras Pampeanas, further supporting the need for targeted sampling to evaluate its genetic structure and potential cryptic diversity.

Climate change adds urgency to this effort, as many birds adjust their elevational ranges to track suitable temperatures (Freeman et al., 2018). Our results indicate that colonization of montane forests occurred repeatedly and independently in different *Pyrrhura* lineages over a relatively short evolutionary timescale. This suggests that Andean environments may provide the thermal gradients necessary to buffer montane-adapted *Pyrrhura* species against rapid warming, whereas Amazonian lowland species, often more restricted in range and dispersal, may be disproportionately vulnerable.

## Acknowledgments

This study was supported by funds granted by NTNU to MDM as part of the Onsager Fellowship Programme, as well as by the Norwegian Directorate for Higher Education and Skills (HKdir) through NORPART award 2021/10475 (“BiGTREE: Biodiversity Genomics Teaching, Research, Exchanges and Education”). Funding was also provided in part by Iridian Genomes grant IRGEN_RG_2021-1345 (“Genomic Studies of Eukaryotic Taxa”) and by the Conselho Nacional de Desenvolvimento Científico e Tecnológico (CNPq, 306989/2023-9), Fundación de Apoyo a la Investigación del Estado de São Paulo (FAPESP, 2013/50297-0) from Brazil, and by the Peder Sather Grant awarded to JC. We thank Letty Salinas (Natural History Museum of San Marcos, Peru) for her continual support and for facilitating access to the valuable collections of the museum, and the curatorial staff of the Institute of Pharmacy and Molecular Biotechnology (IPMB, Germany) for facilitating access to sample material. In addition we want to thank Dr. Martin Päckert for his valued comments on the first manuscript. Special thanks to the Peruvian Government for the research and sampling permit issued by the local authorities (SERFOR N° AUT-IFS-2022-042,). Samples from Peru and Brazil were exported to Norway under CITES permits PE005040/SP, 24BR048160/DF and 14BR012633/DF. Raw sequencing data generated for this study has been deposited on the European Nucleotide Archive under project accession code PRJEB90626.

## Contributions

JML wrote the paper with input from all authors. JML, TH and SP performed molecular procedures on the samples. JML analysed the data with input from MDM and TH. JML, MDM, JC, JB, SH, and CYM contributed to interpreting the results. JB, SH, TH, and MW assisted in sample acquisition and preparation. JML and TH had the original idea for the study. JML and MDM designed the study with input from all authors.

## Notes

### Competing Interest Statement

The authors have declared no competing interest.

