## Supplementary Information for "Phylogenomics and biogeography of the parrot genus *Pyrrhura* with implications for systematics and conservation"

**Table S1.** Systematics of the *Pyrrhura* genus. Current accepted species and subspecies taxa by the South American Classification Committee (Remsen et al., 2025) and the IOC World Bird List (Gill et al., 2025). We also show other described *Pyrrhura* taxa and their corresponding reference.

| Species (SACC) | Species (IOC v15.1) | Subspecies (IOC v15.1) | Other described Taxa |
|---|---|---|---|
| *Pyrrhura cruentata* | *Pyrrhura cruentata* | monotypic | |
| *Pyrrhura devillei* | *Pyrrhura devillei* | monotypic | |
| *Pyrrhura frontalis* | *Pyrrhura frontalis* | *P. frontalis frontalis (=kriegi)* | |
| | | *P. frontalis chiripepe (=borelli)* | |
| *Pyrrhura lepida* | *Pyrrhura lepida* | *P. lepida lepida ( = coerulescens)* | |
| | | *P. lepida coerulescens ( = lepida)* | |
| | | *P. lepida anerythra* | |
| *Pyrrhura perlata* | *Pyrrhura perlata (=rhodogaster)* | monotypic | |
| *Pyrrhura molinae* | *Pyrrhura molinae* | *P. molinae molinae* | |
| | | *P. molinae flavoptera* | |
| | | *P. molinae phoenicura* | |
| | | *P. molinae hypoxantha (=sordida)* | |
| | | *P. molinae restricta* | |

| | | |
|---|---|---|
| | | *P. molinae australis* |
| *Pyrrhura viridicata* | *Pyrrhura viridicata* | monotypic |
| *Pyrrhura egregia* | *Pyrrhura egregia* | *P. egregia egregia* |
| | | *P. egregia obscura* |
| *Pyrrhura melanura* | *Pyrrhura melanura* | *P. melanura melanura* |
| | | *P. melanura berlepschi* |
| | | *P. melanura souancei* |
| | | *P. melanura pacifica* |
| | | *P. melanura chapmani* |
| *Pyrrhura orcesi* | *Pyrrhura orcesi* | monotypic |
| *Pyrrhura rupicola* | *Pyrrhura rupicola* | *P. rupicola rupicola* |
| | | *P. rupicola sandiae* |
| *Pyrrhura albipectus* | *Pyrrhura albipectus* | monotypic |
| *Pyrrhura calliptera* | *Pyrrhura calliptera* | monotypic |
| *Pyrrhura hoematotis* | *Pyrrhura hoematotis* | *P. hoematotis hoematotis* |
| | | *P. hoematotis immarginata* |
| *Pyrrhura rhodocephala* | *Pyrrhura rhodocephala* | monotypic |
| | *Pyrrhura hoffmanni* | *P. hoffmanni hoffmanni* |

| | | | |
|---|---|---|---|
| | | *P. hoffmanni gaudens* | |
| *Pyrrhura pfrimeri* | *Pyrrhura pfrimeri* | monotypic | |
| *Pyrrhura griseipectus* | *Pyrrhura griseipectus* | monotypic | |
| *Pyrrhura leucotis* | *Pyrrhura leucotis* | monotypic | |
| *Pyrrhura picta* | *Pyrrhura picta* | *P. picta picta* | |
| | | *P. picta eisenmanni* | |
| | | *P. picta subandina* | |
| | | *P. picta caeruleiceps* | |
| | | | *P. caeruleiceps pantchenkoi* (Phelps, 1977) |
| | *Pyrrhura emma* | monotypic | |
| | | | *P. emma auricularis* (Zimmer & Phelps, 1949) |
| *Pyrrhura amazonum* | *Pyrrhura amazonum* | *P. amazonum amazonum* | |
| | | | *P. amazonum araguaiaensis* (Arndt & Wink, 2017) |
| | | | *P. amazonum microtera* (Todd, 1947) |
| | | *P. amazonum pallescens (=snethlageae)* | |
| | | *P. amazonum pallescens lucida (=melanoides)* | |

| | | | |
|---|---|---|---|
| | | | *P. pallescens ochrotis* (Miranda-Ribeiro, 1926) |
| *Pyrrhura lucianii* | *Pyrrhura lucianii* | monotypic | |
| | | | *P. lucianii orosaensis* (Arndt & Wink, 2017) |
| *Pyrrhura roseifrons* | *Pyrrhura roseifrons* | *P. roseifrons roseifrons* | |
| | | *P. roseifrons parvifrons* | |
| | | *P. roseifrons peruviana* | |
| | | *P. roseifrons dilutissima* | |
| | | | *P. dilutssima pareneensis* (Arndt & Wink, 2017) |

**Table S2.** The sampling set included in this study. DNA extracts were provided by the Institute of Plant Molecular Biology (IPMB, Germany). In addition, samples were sourced from various museum collections. Taxonomic identification is based on the IOC (v15), and alternative taxonomic identification (not IOC-recognized) is shown within parentheses. Size of the mitochondrial DNA is shown, and incomplete mitogenomes are denoted by an asterisk "*". The last three columns correspond to the raw reads of the whole genome sequencing data, and the mean depth of the nuclear genome and endogenous content after mapping the raw reads to the *Pyrrhura perlata* reference genome (MinQ30).

| Genus | Species IOC | ASTRAL | Subspecies IOC (Additional Taxa ID in shown paranthesis) | Identifier | Year of collection | Country and Locality | Mitogenome length (bp) | NCBI accession code for mitogenome assembly | European Nucleotide Archive (ENA) Study Accession | Raw reads | Mean Sequencing Depth of nuclear genome (MinQ30) | Endogenic Content (MinQ30) | Reference to previous studies |
|---|---|---|---|---|---|---|---|---|---|---|---|---|---|
| Anodorhynchus | hyacinthus | YES | | | | USA: Captive (pet bird) | | | | 59132070 | 9,26 | 0,77 | Hains et al. 2022 (WOUG00000000) |
| Ara | ararauna | YES | | | NA | USA: Captive (pet bird) | | | Data available upon request to Taylor Hains | 88793517 | 13,14 | 0,77 | |
| Aratinga | maculata | YES | | YPM ORN 137433 | 2007 | Suriname | | | | 102185724 | 17,55 | 0,77 | Hains et al. 2022 (JANCNV000000000) |

| | | | | | | | | | | | | | |
|---|---|---|---|---|---|---|---|---|---|---|---|---|---|
| Deroptyus | accipitrinus | YES | | ANSP 188419 | 1997 | Guyana | | | | 121656092 | 22,92 | 0,78 | Hains et al. 2022 (JANHGX000000000) |
| Psittacara | mitratus | YES | | CUMV Bird 52922 | 2005 | Argentina | | | | 67816680 | 12,14 | 0,79 | Hains et al. 2022 (JANEXA000000000) |
| Rhynchopsitta | pachyrhyncha | YES | | | | | | | | 94526110 | 13,07 | 0,68 | Hains et al. 2022 (JANEXL000000000) |
| Pyrrhura | cruentata | YES | | UF 50986 | 2015 | USA: Captive bird (Hill Country Aviary) | 16.985 | ON059971 | PRJNA481551 | 66854138 | 13,79 | 0,85 | Hains et al. 2022 (WUAO00000000) |
| Pyrrhura | cruentata | | | IPMB 35112 | 2005 | Spain: Captive bird, Loro Parque (Ring LP 972172, ExternalNo LV381) | 16.983 | PV917592 | PRJEB90626 | 63444477 | 11,29 | 0,88 | |
| Pyrrhura | egregia | YES | | KU 93445 (Tissue 4090) | 2001 | Guyana: N slope Mount Roraima, E bank Waruma River | 16.993 | ON098133 | PRJNA703578 | 79443067 | 14,14 | 0,84 | Hains et al. 2022 (JANEXG000000000) |
| Pyrrhura | egregia | | | IPMB 35121 | 2005 | Spain: Captive bird, Loro Parque (Ring LP 013197, ExternalNo LV390) | 17.001 | PV917628 | PRJEB90626 | 48289714 | 7,39 | 0,85 | |

| | | | | | | | | | | | | | |
|---|---|---|---|---|---|---|---|---|---|---|---|---|---|
| Pyrrhura | albipectus | YES | | LSUMZ B-6030 | | Ecuador: Morona-Santiago | 16.994 | ON098135 | PRJNA703575 | 75189912 | 11,91 | 0,77 | Hains et al. 2022 (JANFOL000000000) |
| Pyrrhura | calliptera | | | MTDC-58653 | NA | Colombia | 16.993 | PV917629 | PRJEB90626 | 14902375 | 0,86 | 0,65 | |
| Pyrrhura | calliptera | | | MTDC-58652 | NA | Colombia | 16.993 | PV917630 | PRJEB90626 | 43085246 | 2,25 | 0,63 | |
| Pyrrhura | calliptera | YES | | FMNH 53678 | | Colombia: Bogota | 16.993 | | | 239409566 | 10,21 | 0,68 | |
| Pyrrhura | hoematotis | YES | | UMMZ 589 | 1989 | Venezuela: Distrito Federal, Highway between "Colonia Tovar" (Aragua) & Chichiriviche,La Fundacion | 16.993 | ON122997 | PRJNA491344 | 127036247 | 18,20 | 0,73 | Hains et al. 2022 (JANFOO000000000) |
| Pyrrhura | melanura | | cf. souancei (based on morphology) | IPMB 35146 | 2005 | Spain: Captive bird, Loro Parque (Ring 055106, ExternalNo F12) | 16.993 | PV917631 | PRJEB90626 | 48733894 | 8,66 | 0,87 | |
| Pyrrhura | melanura | YES | melanura | FMNH 456474 | 2007 | Brazil: Amazonas, Maraa, Lago Cumapi | 16.993 | ON123000 | PRJNA703582 | 77274700 | 11,84 | 0,77 | Hains et al. 2022 (JANFOK000000000) |

| | | | | | | | | | | | | | |
|---|---|---|---|---|---|---|---|---|---|---|---|---|---|
| Pyrrhura | melanura | | cf. berlepschi (based on morphology) | IPMB 65832 | 2012 | | 16.992 | PV917632 | PRJEB90626 | 57886610 | 8,11 | 0,83 | |
| Pyrrhura | melanura | YES | berlepschi | ANSP 176701 | 1984 | Ecuador: Morona-Santiago, E of Logrono; W slope of Cordillera de Cutucu | 16.994 | | Data available upon request to Taylor Hains | 66973502 | 7,75 | 0,81 | |
| Pyrrhura | melanura | YES | chapmani | FMNH 281936 | 1967 | Colombia: Putumayo, Mocoa | 16.993 | OR209191 | PRJNA807740 | 133423493 | 13,93 | 0,77 | Hains et al. 2022 (JANZXS000000000) |
| Pyrrhura | melanura | | cf. pacifica (based on morphology) | IPMB 35145 | 2005 | Spain: Captive bird, Loro Parque (Ring LP013337, External No LV1104) | 16.997 | PV917634 | PRJEB90626 | 42307015 | 7,40 | 0,87 | |
| Pyrrhura | melanura | | cf. melanura (based on morphology) | IPMB 35310 | 2007 | Spain: Captive bird, Loro Parque (ExternalNo U289) | 16.997 | PV917633 | PRJEB90626 | 30156299 | 2,30 | 0,69 | |
| Pyrrhura | orcesi | YES | | LSUMZ B-7803 | NA | Ecuador: El Oro Province | 16.990 | ON123002 | PRJNA703583 | 79256599 | 13,37 | 0,80 | Hains et al. 2022 (JANHHA000000000) |

| | | | | | | | | | | | | | |
|---|---|---|---|---|---|---|---|---|---|---|---|---|---|
| Pyrrhura | rhodocephala | | | FMNH 53433 | 1920 | Venezuela: Merida, Rio Mucujun | 16.992 | | Data available upon request to Taylor Hains | 51966950 | 2,45 | 0,36 | |
| Pyrrhura | rhodocephala | YES | | IPMB 35167 | 2005 | Spain: Captive bird, Loro Parque (Ring CVD96023, ExternalNo LV366) | 16.991 | PV917639 | PRJEB90626 | 38866513 | 7,24 | 0,84 | |
| Pyrrhura | hoffmanni | YES | cf. gaudens | LSUMZ B-28202 | NA | Panama: Chiriqui Province | 16.994 | ON122998 | PRJNA703579 | 70183969 | 11,30 | 0,80 | Hains et al. 2022 (JANHGK000000000) |
| Pyrrhura | hoffmanni | | cf. gaudens | IPMB 35130 | 2005 | Spain: Captive bird, Loro Parque (Ring LP 003061, ExternalNo LV369) | 16.995 | PV917638 | PRJEB90626 | 27742093 | 5,35 | 0,85 | |
| Pyrrhura | viridicata | YES | | FMNH 72327 | 1926 | Colombia: Magdalena, La Cumbre, Sierra Nevada de Santa Marta | 16.992 | | Data available upon request to Taylor Hains | 43903917 | 2,58 | 0,42 | |
| Pyrrhura | frontalis | YES | | KU 86335 (Tissue 1433) | NA | Unkown: Wild bird confiscated by USFWS | 16.987 | | Data available upon request to Taylor Hains | 479619652 | 26,82 | 0,75 | |

| | | | | | | | | | | | | | |
|---|---|---|---|---|---|---|---|---|---|---|---|---|---|
| Pyrrhura | frontalis | | cf. chiripepe | LSUMZ B-25884 | NA | Paraguay: Caazapá Department | 16.991 | ON122995 | PRJNA481541 | 68654309 | 13,22 | 0,80 | Hains et al. 2022 (JAAAKN000000000) |
| Pyrrhura | frontalis | | cf. chiripepe | LGEMA 18832 | | Brazil: RS, Sao Francisco de Paula, Parque Municipal da RONDA | 16.992 | PV917635 | PRJEB90626 | 55490779 | 6,98 | 0,78 | |
| Pyrrhura | frontalis | | | IPMB 35126 | 2005 | Spain: Captive bird, Loro Parque (Ring LP 993057, ExternalNo LV1090) | 16931* | PV917645 | PRJEB90626 | 57877542 | 6,33 | 0,83 | |
| Pyrrhura | molinae | | | | NA | USA: Captive, pet store from New York (Name: Hera) | 16.991 | | PRJNA481548 | 67262669 | 14,67 | 0,87 | Hains et al. 2022 (JAADKS000000000) |
| Pyrrhura | molinae | | cf. restricta | IPMB 35149 | 2005 | Spain: Captive bird, Loro Parque (ExternalNo Q251) | 16.992 | PV917636 | PRJEB90626 | 55230737 | 8,68 | 0,83 | |
| Pyrrhura | molinae | | | IPMB 35148 | 2005 | Spain: Captive bird, Loro Parque (Ring CO0864, ExternalNo LV1103) | 16448* | PV917647 | PRJEB90626 | 10788362 | 1,92 | 0,85 | |
| Pyrrhura | molinae | YES | cf. hypoxantha | KU 90195 | 1999 | Paraguay: Alto Paraguay, Picada Chovoreca | 16.993 | | Data available upon request to Taylor Hains | 131942481 | 17,13 | 0,78 | |

| | | | | | | | | | | | | | |
|---|---|---|---|---|---|---|---|---|---|---|---|---|---|
| Pyrrhura | devillei | YES | | MCZ 154556 | 1930 | Brazil: Mato Grosso do Sul | 16.992 | | PRJNA491342 | 75245962 | 4,26 | 0,34 | |
| Pyrrhura | lepida | YES | coerulescens | FMNH GUR 296 | NA | Brazil: Gurupi, State of Tocantis | 16.993 | | PRJNA481549 | 75816374 | 11,81 | 0,81 | Hains et al. 2022 (JAABNU000000000) |
| Pyrrhura | lepida | YES | coerulescens | FMNH GUR 300 | NA | Brazil: Gurupi, State of Tocantis | 16.992 | | Data available upon request to Taylor Hains | 234516777 | 22,17 | 0,84 | |
| Pyrrhura | lepida | | lepida | IPMB 35135 | 2005 | Spain: Captive bird, Loro Parque (Ring 055007, ExternalNo F11) | 16.993 | PV917640 | PRJEB90626 | 36286331 | 6,74 | 0,85 | |
| Pyrrhura | lepida | | lepida | IPMB 35311 | 2007 | Spain: Captive bird, Loro Parque (ExternalNo U277) | 16.993 | PV917641 | PRJEB90626 | 49405937 | 0,68 | 0,52 | |
| Pyrrhura | perlata | YES | | FMNH 389699 | 1986 | Brazil: Rondonia, Cachoeira Nazare, W bank Rio Jiparana | 16.994 | | Data available upon request to Taylor Hains | 398357959 | 31,54 | 0,85 | |
| Pyrrhura | perlata | | | | NA | USA: Captive, pet store from New York (Name: Kira) | 16.993 | | PRJNA481550 | 66853523 | 13,59 | 0,83 | Hains et al. 2022 (JAAGVT000000000) |

| | | | | | | | | | | | | | |
|---|---|---|---|---|---|---|---|---|---|---|---|---|---|
| Pyrrhura | perlata | | | IPMB 35155 | 2005 | Spain: Captive bird, Loro Parque (Ring CO2903, ExternalNo LV377) | 16.993 | PV917637 | PRJEB90626 | 23209976 | 4,31 | 0,84 | |
| Pyrrhura | rupicola | | cf. rupicola | IPMB 35175 | 2005 | Spain: Captive bird, Loro Parque (Ring AFE084, ExternalNo LV384) | 16.993 | PV917643 | PRJEB90626 | 29964453 | 5,84 | 0,87 | |
| Pyrrhura | rupicola | | cf. sandiae | IPMB 35176 | 2005 | Spain: Captive bird, Loro Parque (Ring LP962139, ExternalNo LV360) | 16.978 | PV917642 | PRJEB90626 | 34505092 | 6,28 | 0,84 | |
| Pyrrhura | rupicola | YES | rupicola | LSUMZ B-10554 | NA | Peru: Ucayali | 16.993 | | PRJNA703587 | 93795507 | 16,09 | 0,78 | Hains et al. 2022 (JANFHV000000000) |
| Pyrrhura | emma | | | IPMB 65038 | 2012 | Spain: Captive bird, Loro Parque (966501009) | 16.988 | PV917627 | PRJEB90626 | 16781063 | 1,82 | 0,78 | Arndt & Wink 2017 |
| Pyrrhura | emma | YES | (auricularis) | FMNH 97206 | 1932 | Venezuela: Sucre, Mt Turumiquire | 16.993 | | Data available upon request to Taylor Hains | 52454199 | 3,48 | 0,49 | |
| Pyrrhura | emma | | emma (♂) x amazonum (♀) | IPMB 35139 | 2005 | Spain: Captive bird, Loro Parque (Ring 2700031, ExternalNo LV385) | 16.990 | PV917614 | PRJEB90626 | 45447879 | 4,13 | 0,83 | |

| | | | | | | | | | | | | | |
|---|---|---|---|---|---|---|---|---|---|---|---|---|---|
| Pyrrhura | picta | | picta | IPMB 35163 | 2005 | Spain: Captive bird, Loro Parque (Ring LP982136, ExternalNo LV375) | 16.978 | PV917613 | PRJEB90626 | 28850040 | 5,03 | 0,81 | |
| Pyrrhura | picta | YES | picta | FMNH 395728 | 1992 | Brazil: Roraima, Vila Surumu, 7 km W, Rio Surumu | 16.979 | ON123004 | PRJNA703585 | 70916653 | 10,15 | 0,77 | Hains et al. 2022 (JANHHB000000000) |
| Pyrrhura | picta | YES | eisenmanni | ANSP 189116 | 1996 | Panama: Veraguas, Cascajillaso | 16.993 | | Data available upon request to Taylor Hains | 75264496 | 11,13 | 0,77 | |
| Pyrrhura | picta | YES | subandina | ANSP 160666 | 1949 | Colombia: Bolivar, Cerro Murrucucu | 16.980 | | Data available upon request to Taylor Hains | 53955881 | 5,35 | 0,78 | |
| Pyrrhura | leucotis | | | VM-NTNU 4623 | 1960 | Brazil | 16.985 | PV917622 | PRJEB90626 | 22977078 | 1,04 | 0,69 | |
| Pyrrhura | leucotis | YES | | IPMB 35142 | 2005 | Spain: Captive bird, Loro Parque (Ring PPGC 019, ExternalNo LV378) | 16.984 | PV917621 | PRJEB90626 | 57719279 | 7,71 | 0,86 | |

| | | | | | | | | | | | | | |
|---|---|---|---|---|---|---|---|---|---|---|---|---|---|
| Pyrrhura | leucotis | | | IPMB 60411 | 2011 | Captive Bird (ExternalNo 189) | 16.983 | PV917620 | PRJEB90626 | 20239635 | 3,30 | 0,83 | Arndt & Wink 2017 |
| Pyrrhura | leucotis | | leucotis (♂) x griseipectus (♀) | IPMB 60410 | 2011 | Captive Bird (ExternalNo 183) | 16.983 | PV917626 | PRJEB90626 | 13262542 | 2,53 | 0,85 | Arndt & Wink 2017 |
| Pyrrhura | griseipectus | | | IPMB 60412 | 2011 | Captive Bird (ExternalNo 740) | 16.983 | PV917623 | PRJEB90626 | 36496444 | 2,40 | 0,85 | Arndt & Wink 2017 |
| Pyrrhura | griseipectus | | | IPMB 60413 | 2011 | Captive Bird | 16.983 | PV917624 | PRJEB90626 | 20995804 | 3,06 | 0,83 | Arndt & Wink 2017 |
| Pyrrhura | griseipectus | | | IPMB 35141 | 2005 | Spain: Captive bird, Loro Parque (Ring AZ03667P04001, ExternalNo LV1101) | 16.984 | PV917625 | PRJEB90626 | 13159741 | 2,43 | 0,85 | |
| Pyrrhura | griseipectus | YES | | UF 50988 | NA | USA: Captive bird (Hill Country Aviary) | 16.986 | ON122996 | PRJNA598107 | 86066530 | 17,91 | 0,86 | Hains et al. 2022 (JAAAKS000000000) |
| Pyrrhura | amazonum | YES | pallescens | FMNH 389694 | 1986 | Brazil: Rondonia, Cachoeira Nazare, W bank Rio Jiparana | 16.991 | ON098132 | PRJNA703576 | 79743483 | 13,22 | 0,83 | Hains et al. 2022 (JANFON000000000) |

| | | | | | | | | | | | | | |
|---|---|---|---|---|---|---|---|---|---|---|---|---|---|
| Pyrrhura | amazonum | | amazonum (microtera) | IPMB 77902 (MZUSP) | NA | Brazil: Para, Altamira, east of Rio Xingu | 16.989 | PV917619 | PRJEB90626 | 18843766 | 2,66 | 0,78 | Arndt & Wink 2017 |
| Pyrrhura | amazonum | | amazonum (microtera) | IPMB 77903 (MZUSP) | NA | Brazil: Para, Altamira, east of Rio Xingu | 16.990 | PV917616 | PRJEB90626 | 12027650 | 1,85 | 0,81 | Arndt & Wink 2017 |
| Pyrrhura | amazonum | YES | amazonum (araguaiensis) | IPMB 88892 (MZUSP) | NA | Brazil: Santana do Araguaia, Faz. Fartura | 16.990 | PV917615 | PRJEB90626 | 55417008 | 7,51 | 0,82 | Arndt & Wink 2017 |
| Pyrrhura | pfrimeri | YES | | FMNH 75204 | 1930 | Brazil: Goias, Nova Roma, Rio Parana | 16.976 | | Data available upon request to Taylor Hains | 52259770 | 4,42 | 0,57 | |
| Pyrrhura | pfrimeri | | | IPMB 60414 | 2011 | Captive Bird (ExternalNo 1032) | 16.976 | PV917593 | PRJEB90626 | 70106866 | 10,79 | 0,83 | Arndt & Wink 2017 |
| Pyrrhura | pfrimeri | | | IPMB 60416 | 2011 | Captive Bird | 16.977 | PV917594 | PRJEB90626 | 57156299 | 9,19 | 0,84 | Arndt & Wink 2017 |
| Pyrrhura | roseifrons | | dilutissima (dilutissima) | IPMB 79372 | 2014 | Peru: Ayacucho, Pumorini, Kimbiri Alto | 16.994 | PV917610 | PRJEB90626 | 50041533 | 7,22 | 0,85 | Arndt & Wink 2017 |

| | | | | | | | | | | | | | |
|---|---|---|---|---|---|---|---|---|---|---|---|---|---|
| Pyrrhura | roseifrons | | dilutissima (paraneensis) | MUSM 26713 (Type) | NA | Peru: Junin, Sondobeni (Rio Tambo) | 16.997 | PV917612 | PRJEB90626 | 16078354 | 1,33 | 0,77 | |
| Pyrrhura | roseifrons | | dilutissima (paraneensis) | IPMB 76252 | 2013 | Peru: Junin, Sondobeni (Rio Tambo) | 16996* | PV917609 | PRJEB90626 | 13251668 | 2,38 | 0,86 | Arndt & Wink 2017 |
| Pyrrhura | roseifrons | YES | dilutissima (paraneensis) | IPMB 76253 | 2013 | Peru: Junin, Sondobeni (Rio Tambo) | 16.996 | PV917611 | PRJEB90626 | 43020270 | 7,54 | 0,86 | Arndt & Wink 2017 |
| Pyrrhura | roseifrons | YES | | LSUMZ 206368 | 2011 | Peru: Ucayali: north ridge Quebrada Quirapokiari, 22.87 km SW mouth Rio Cohengua | 16.992 | | | 25162606 | 3,06 | 0,81 | |
| Pyrrhura | roseifrons | | | IPMB 35164 | 2005 | Spain: Captive bird, Loro Parque (Ring LP982136, ExternalNo LV375) | 16.992 | PV917599 | PRJEB90626 | 24282204 | 4,52 | 0,84 | |
| Pyrrhura | roseifrons | | (roseifrons II) | IPMB 65039 | 2012 | Spain: Captive bird, Loro Parque (003017) | 16.992 | PV917598 | PRJEB90626 | 52868244 | 8,19 | 0,84 | Arndt & Wink 2017 |
| Pyrrhura | roseifrons | | (roseifrons II) | IPMB 77887 | NA | Brazil: Captive bird (Pedro Teles, Brasilien) Criadouri Boa esperanca (SDB196), probably rio Juruá | 16.992 | PV917600 | PRJEB90626 | 42914779 | 6,80 | 0,82 | Arndt & Wink 2017 |

| | | | | | | | | | | | | | |
|---|---|---|---|---|---|---|---|---|---|---|---|---|---|
| Pyrrhura | roseifrons | YES | parvifrons | IPMB 79375 | 2014 | Peru: "La Carachamera", altura km 38 de la carretera Tarapoto a Yurimagua, San Martin | 16.981 | PV917602 | PRJEB90626 | 69190489 | 8,66 | 0,86 | Arndt & Wink 2017 |
| Pyrrhura | roseifrons | YES | peruviana | MVZ 165115 | NA | Peru: Entsa [=creek] Ketai, Rio Cenepa, Amazonas | 16.991 | ON123003 | PRJNA703581 | 123590848 | 20,25 | 0,84 | Hains et al. 2022 (JANIAF000000000) |
| Pyrrhura | roseifrons | | peruviana | IPMB 76251 | 2013 | Peru: Huampami, Rio Cenepa, Amazonas | 16979 (r) | PV917603 | PRJEB90626 | 46581648 | 3,76 | 0,79 | Arndt & Wink 2017 |
| Pyrrhura | lucianii | YES | | IPMB 77896 (MZUSP J372) | NA | Brazil: Rio Madeira, Abuna | 16980 (r) | PV917604 | PRJEB90626 | 74020131 | 9,94 | 0,81 | Arndt & Wink 2017 |
| Pyrrhura | lucianii | | | IPMB 77897 (MZUSP J1126) | NA | Brazil: Rio Madeira, Abuna, Barreiro | 16.990 | PV917605 | PRJEB90626 | 20474758 | 3,09 | 0,83 | Arndt & Wink 2017 |
| Pyrrhura | lucianii | | (orosaensis) | MUSM 32474 | 2012 | Peru: Rio Orosa, near a small collpa | 16979 (r) | PV917608 | PRJEB90626 | 14240266 | 1,35 | 0,77 | |
| Pyrrhura | lucianii | | (orosaensis) | MUSM 32475 | 2012 | Peru: Rio Orosa, near a small collpa | 16.989 | PV917607 | PRJEB90626 | 18767565 | 1,69 | 0,78 | |

**Table S3**. Inter- and intra-specific genetic distances within *Pyrrhura*. Mean pairwise genetic distances are determined from the branch lengths in the mitochondrial genome phylogeny. Genetic distances are measured in average pairwise tree distance.

| **Species** | **Closest matching species** | **Intra-specific genetic distance** | **Inter-specific genetic distance to closest relative** |
|---|---|---|---|
| *P. cruentata* | *P. l. coerulescens* | 0.002 | 0.074 |
| *P. devillei* | *P. frontalis* | **NA** | 0.009 |
| *P. frontalis* | *P. devillei* | 0.001 | 0.009 |
| *P. lepida coerulescens* | *P. rupicola* | 0.002 | 0.027 |
| *P. perlata* | *P. molinae* | **NA** | 0.003 |
| *P. perlata* (captive clade) | *P. molinae* | 1.22E-4 | 0.012 |
| *P. molinae* | *P. perlata* | **NA** | 0.003 |
| *P. molinae* (captive clade) | *P. perlata* (captive clade) | 0.003 | 0.019 |
| *P. viridicata* | *P. rhodocephala* | **NA** | 0.014 |
| *P. egregia* | *P. albipectus* | 0.015 | 0.018 |
| *P. melanura* | *P. m. chapmani* | **NA** | 0.007 |
| *P. melanura berlepschi* | *P. albipectus* | **NA** | 6.10E-5 |
| *P. melanura chapmani* | *P. hoematotis* | **NA** | 0.006 |
| *P. melanura* (captive clade) | *P. m. chapmani* | 0.001 | 0.009 |
| *P. melanura souancei* (captive individual) | *P. albipectus* | **NA** | 0.002 |
| *P. orcesi* | *P. hoffmanni* | **NA** | 0.021 |
| *P. rupicola* | *P. l. coerulescens* | 0.007 | 0.027 |
| *P. albipectus* | *P. m. berlepschi* | **NA** | 6.10E-5 |
| *P. calliptera* | *P. albipectus* | 0.002 | 0.003 |
| *P. hoematotis* | *P. m. chapmani* | **NA** | 0.006 |
| *P. rhodocephala* | *P. viridicata* | 8.37E-4 | 0.014 |
| *P. hoffmanni* | *P. rhodocephala* | 4.83E-4 | 0.018 |
| *P. pfrimeri* | *P. r. dilutissima* | 0.001 | 0.021 |
| *P. griseipectus* | *P. leucotis* | 0.005 | 0.011 |
| *P. leucotis* | *P. griseipectus* | 0.003 | 0.011 |
| *P. picta picta* | *P. p. emma* | 0.001 | 0.018 |
| *P. picta eisenmanni* | *P. p. subandina* | **NA** | 0.003 |
| *P. picta subandina* | *P. p. eisenmanni* | **NA** | 0.003 |
| *P. emma* | *P. p. picta* | 0.013 | 0.018 |
| *P. amazonum/P. a. pallescens* | *P. leucotis* | 0.003 | 0.012 |
| *P. lucianii lucianii* | *P. l. orosaensis* | 0.002 | 0.004 |
| *P. lucianii orosaensis* | *P. l. lucianii* | 0.002 | 0.004 |
| *P. roseifrons roseifrons* | *P. r. dilutissima* | 3.35E-4 | 0.012 |
| *P. roseifrons peruviana* | *P. r. parvifrons* | 0.001 | 0.003 |
| *P. roseifrons parvifrons* | *P. r. peruviana* | **NA** | 0.003 |
| *P. roseifrons dilutissima* | *P. r. peruviana* | 0.001 | 0.011 |

**Table S4.** Identification and annotating tRNA genes in the mitochondrial genome of *P. r. parvifrons* MUSM 32747 using tRNA-scan-SE v2.0 (Lowe & Chan, 2016).

| tRNA # | tRNA Begin | tRNA End | tRNA Type | Anticodon | Intron Begin | Intron End | Infernal Score | Note |
|---|---|---|---|---|---|---|---|---|
| 1 | 1 | 67 | Phe | GAA | 0 | 0 | 70.5 | None |
| 2 | 1039 | 1110 | Val | TAC | 0 | 0 | 86.6 | None |
| 3 | 2671 | 2745 | Leu | TAA | 0 | 0 | 107.9 | None |
| 4 | 3731 | 3802 | Ile | GAT | 0 | 0 | 89.9 | None |
| 5 | 3878 | 3946 | Met | CAT | 0 | 0 | 104.4 | None |
| 6 | 4987 | 5057 | Trp | TCA | 0 | 0 | 106.2 | None |
| 7 | 6971 | 7039 | Asp | GTC | 0 | 0 | 79.4 | None |
| 8 | 7727 | 7795 | Lys | TTT | 0 | 0 | 83.8 | None |
| 9 | 9422 | 9490 | Gly | TCC | 0 | 0 | 77.0 | None |
| 10 | 9842 | 9911 | Arg | TCG | 0 | 0 | 87.7 | None |
| 11 | 11596 | 11664 | His | GTG | 0 | 0 | 93.6 | None |
| 12 | 11665 | 11730 | Ser | GCT | 0 | 0 | 55.6 | No D-arm |
| 13 | 11730 | 11800 | Leu | TAG | 0 | 0 | 109.2 | None |
| 14 | 14769 | 14836 | Thr | TGT | 0 | 0 | 73.0 | None |
| 15 | 15497 | 15428 | Glu | TTC | 0 | 0 | 87.3 | None |
| 16 | 14909 | 14841 | Pro | TGG | 0 | 0 | 86.4 | None |

| | | | | | | | | |
|---|---|---|---|---|---|---|---|---|
| 17 | 6967 | 6891 | Ser | TGA | 0 | 0 | 92.5 | None |
| 18 | 5342 | 5273 | Tyr | GTA | 0 | 0 | 96.7 | None |
| 19 | 5272 | 5206 | Cys | GCA | 0 | 0 | 56.6 | None |
| 20 | 5203 | 5129 | Asn | GTT | 0 | 0 | 86.7 | None |
| 21 | 5127 | 5059 | Ala | TGC | 0 | 0 | 88.8 | None |
| 22 | 3878 | 3808 | Gln | TTG | 0 | 0 | 93.2 | None |

**Table S5.** *De-novo* annotation of the mitochondrial genome of *P. roseifrons parvifrons* (MUSM 32747) using MITOS 2 within Galaxy v2.1.9 (https://usegalaxy.org).

| Start | End | Gene | Direction |
|---|---|---|---|
| 0 | 67 | trnF(gaa) | + |
| 66 | 1039 | rrnS | + |
| 1038 | 1110 | trnV(tac) | + |
| 1112 | 2670 | rrnL | + |
| 2670 | 2745 | trnL2(taa) | + |
| 2751 | 3725 | nad1 | + |
| 3730 | 3802 | trnI(gat) | + |
| 3807 | 3878 | trnQ(ttg) | - |
| 3877 | 3946 | trnM(cat) | + |
| 3946 | 4987 | nad2 | + |
| 4986 | 5057 | trnW(tca) | + |
| 5058 | 5127 | trnA(tgc) | - |
| 5128 | 5203 | trnN(gtt) | - |
| 5205 | 5272 | trnC(gca) | - |
| 5271 | 5342 | trnY(gta) | - |
| 5351 | 6899 | cox1 | + |
| 6890 | 6967 | trnS2(tga) | - |
| 6970 | 7039 | trnD(gtc) | + |
| 7041 | 7725 | cox2 | + |
| 7726 | 7795 | trnK(ttt) | + |
| 7796 | 7964 | atp8 | + |
| 7954 | 8638 | atp6_0 | + |

| | | | |
|---|---|---|---|
| 8633 | 8687 | atp6_1 | + |
| 8637 | 9421 | cox3 | + |
| 9421 | 9490 | trnG(tcc) | + |
| 9490 | 9676 | nad3_1 | + |
| 9665 | 9839 | nad3_0 | + |
| 9841 | 9911 | trnR(tcg) | + |
| 9912 | 10209 | nad4l | + |
| 10202 | 11580 | nad4 | + |
| 11595 | 11664 | trnH(gtg) | + |
| 11664 | 11730 | trnS1(gct) | + |
| 11729 | 11800 | trnL1(tag) | + |
| 11779 | 13615 | nad5 | + |
| 13628 | 14768 | cob | + |
| 14768 | 14836 | trnT(tgt) | + |
| 14840 | 14909 | trnP(tgg) | - |
| 14912 | 15425 | nad6 | - |
| 15427 | 15497 | trnE(ttc) | - |

**Table S6.** Partitions and nucleotide substitution models used for the mitogenome phylogenetic analysis. Model selection (under BIC), best partition scheme, and phylogenetic tree construction were performed in IQ-TREE v2.0.3 (Nguyen et al., 2015) using the following command: iqtree -s pyrrhura_mit.phy -p pyrrhura_partition.nex -m MFP+MERGE -B 1000 -nt AUTO. Initially, the mitogenome was partitioned by gene and codon, resulting in 67 schemes. The algorithm merged most of them as shown in the following table:

| **Partition** | **Genes** | **Model** | **Description** |
|---|---|---|---|
| 1 | 12S rRNA, 16S rRNA, tRNA-Val, tRNA-Phe, tRNA-Glu, ND1-1, ND2-1, tRNA-Lys, ATP8-1, ATP6-1, ND4-1, ND5-1, ND6-1, ATP8-2, ND6-2 | TPM2+F+I+G4 | All rRNA, tRNAs, first and second codon of PCGs |
| 2 | tRNA-Leu1, tRNA-Ile, tRNA-Gln, tRNA-Met, tRNA-Trp, tRNA-Ala, tRNA-Asn, tRNA-Cys, tRNA-Tyr, tRNA-Ser1, tRNA-Asp, tRNA-Gly, tRNA-Arg, tRNA-His, tRNA-Ser2, tRNA-Leu2, tRNA-Thr, tRNA-Pro, COX1-1, COX3-1, ND4L-1, CytB-1, COX2-1, ND3-1, | TPM2+F+I+G4 | most tRNAs, first codon of PCGs |
| 3 | ND1-2, ND2-2, COX1-2, COX2-2, ATP6-2, COX3-2, ND3-2, ND4L-2, ND4-2, ND5-2, CytB-2 | TIM3+F+I+G4 | Second codon of most PCGs |
| 4 | ND1-3, ND2-3, COX1-3, COX2-3, ATP8-3, ATP6-3, COX3-3, ND3-3, ND4L-3, ND4-3, ND5-3, CytB-3, ND6-3 | TIM3+F+R3 | Third codon of all PCGs |
| 5 | Control Region | TPM2+F+R3 | Control region |

**Table S7**. Five discrete areas representing different South American biogeographical regions based on Morrone et al. (2022) were utilized to infer the ancestral geographical ranges of the genus *Pyrrhura*.

| **CODE** | **Major Biomes** | **Regions (Morrone et al. 2022)** |
|---|---|---|
| NSM | North South America | Mosquito, Magdalena, Guatuso-Talamanca, Guajira, Western Ecuador, Choco-Darien, Venezuelan, Sabana, Puntarenas-Chiriqui |
| NAM | North Amazonia | Ucayali, Napo, Imeri, Roraima, Guianan, Guianan Lowlands |
| SAM | South Amazonia | Yungas, Amazonia |
| DDI | Dry Diagonal | Chaco, Cerrado, Caatinga |
| ATF | Atlantic Forest | Southern Espinhaco, Atlantic, Parana Forest, Araucaria Forest |

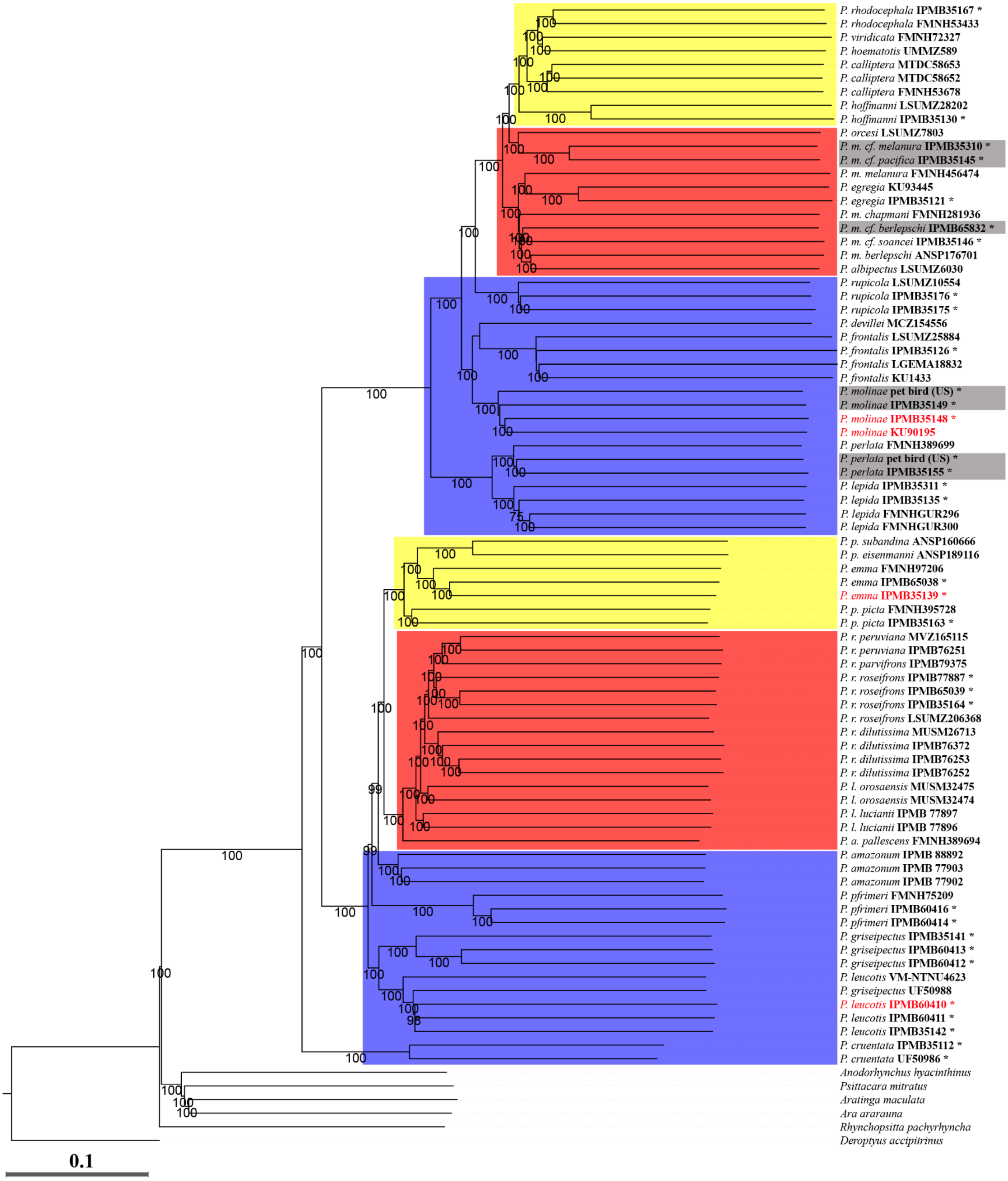
P. rhodocephala IPMB35167 *
P. rhodocephala FMNH53433
P. viridicata FMNH72327
P. hoematotis UMMZ589
P. calliptera MTDC58653
P. calliptera MTDC58652
P. calliptera FMNH53678
P. hoffmanni LSUMZ28202
P. hoffmanni IPMB35130 *
P. orcesi LSUMZ7803
P. m. cf. melanura IPMB35310 *
P. m. cf. pacifica IPMB35145 *
P. m. melanura FMNH456474
P. egregia KU93445
P. egregia IPMB35121 *
P. m. chapmani FMNH281936
P. m. cf. berlepschi IPMB65832 *
P. m. cf. soancei IPMB35146 *
P. m. berlepschi ANSP176701
P. albipectus LSUMZ6030
P. rupicola LSUMZ10554
P. rupicola IPMB35176 *
P. rupicola IPMB35175 *
P. devillei MCZ154556
P. frontalis LSUMZ25884
P. frontalis IPMB35126 *
P. frontalis LGEMA18832
P. frontalis KU1433
P. molinae pet bird (US) *
P. molinae IPMB35149 *
P. molinae IPMB35148 *
P. molinae KU90195
P. perlata FMNH389699
P. perlata pet bird (US) *
P. perlata IPMB35155 *
P. lepida IPMB35311 *
P. lepida IPMB35135 *
P. lepida FMNHGUR296
P. lepida FMNHGUR300
P. p. subandina ANSP160666
P. p. eisenmanni ANSP189116
P. emma FMNH97206
P. emma IPMB65038 *
P. emma IPMB35139 *
P. p. picta FMNH395728
P. p. picta IPMB35163 *
P. r. peruviana MVZ165115
P. r. peruviana IPMB76251
P. r. parvifrons IPMB79375
P. r. roseifrons IPMB77887 *
P. r. roseifrons IPMB65039 *
P. r. roseifrons IPMB35164 *
P. r. roseifrons LSUMZ206368
P. r. dilutissima MUSM26713
P. r. dilutissima IPMB76372
P. r. dilutissima IPMB76253
P. r. dilutissima IPMB76252
P. l. orosaensis MUSM32475
P. l. orosaensis MUSM32474
P. l. luciani IPMB 77897
P. l. luciani IPMB 77896
P. a. pallescens FMNH389694
P. amazonum IPMB 88892
P. amazonum IPMB 77903
P. amazonum IPMB 77902
P. pfrimeri FMNH75209
P. pfrimeri IPMB60416 *
P. pfrimeri IPMB60414 *
P. griseipectus IPMB35141 *
P. griseipectus IPMB60413 *
P. griseipectus IPMB60412 *
P. leucotis VM-NTNU4623
P. griseipectus UF50988
P. leucotis IPMB60410 *
P. leucotis IPMB60411 *
P. leucotis IPMB35142 *
P. cruentata IPMB35112 *
P. cruentata UF50986 *
Anodorhynchus hyacinthinus
Psittacara mitratus
Aratinga maculata
Ara ararauna
Rhynchopsitta pachyrhyncha
Deroptyus accipitrinus
0.1

**Figure S1.** Distance-based phylogenetic tree based on low-depth Whole Genome Sequencing data. Distances were estimated using NGSdist (Vieira et al., 2016), which can handle the uncertainty of genotype assignment by utilizing genotype likelihoods (GL) directly. Samples with red labels correspond to the unexpected placements, and the grey background corresponds to the captive clades observed in the mitogenome-based tree (Figure 2). Asterisks (*) on the label name indicate samples with captive origin. Yellow, red and blue backgrounds on the branches correspond to the topology observed in the phylogeny estimated using ASTRAL (Figure 1). The scale bar corresponds to the calculated genetic distance.

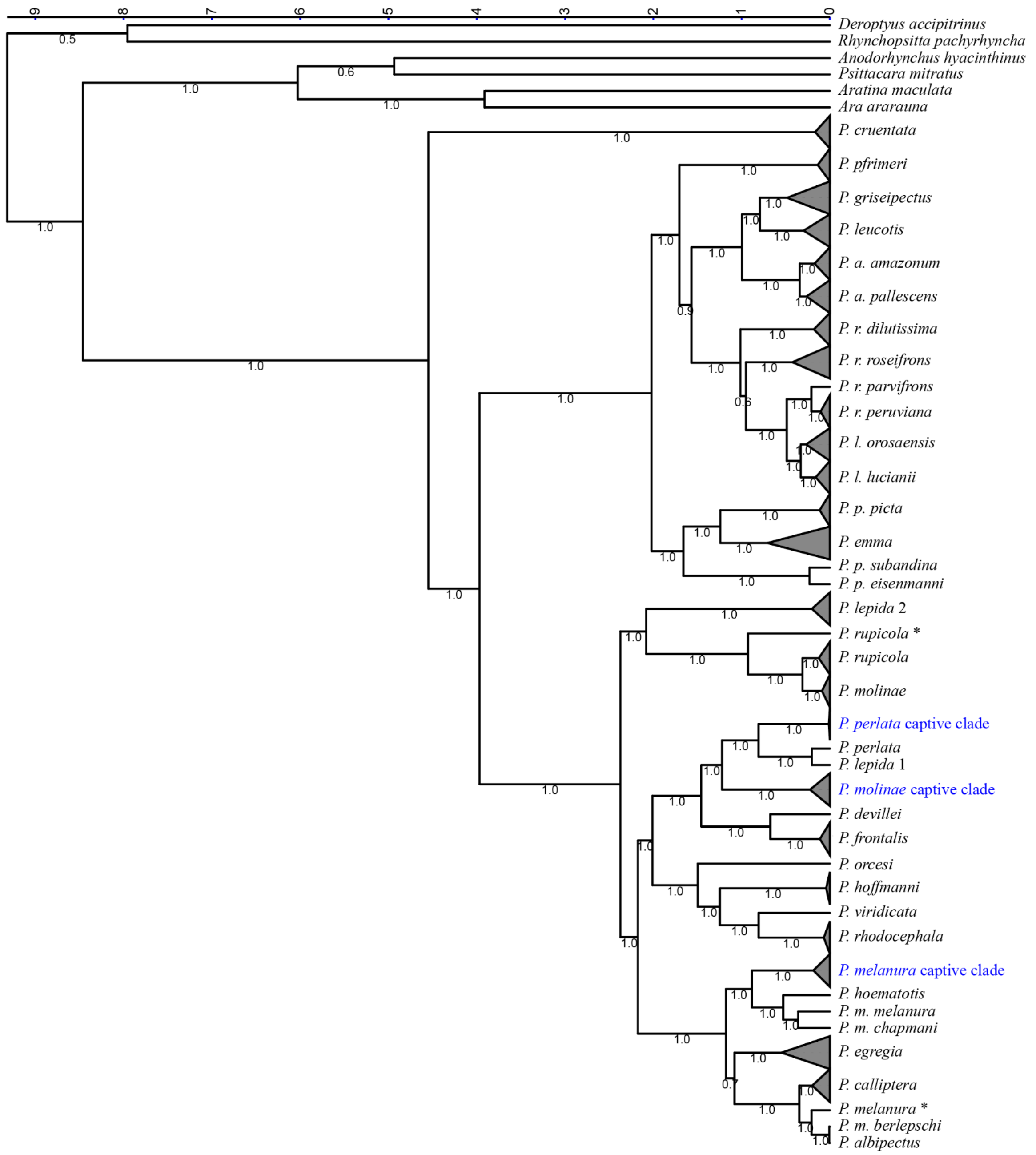

**Figure S2.** Time-calibrated Bayesian phylogeny based on mitochondrial genomes, estimated using BEAST 2.7. Monophyletic clades has been collapsed, and all of them except for the captive clades (in blue) include at least one wild specimen. Divergent captive samples are noted with an asterisk. Posterior probabilities are shown on branches. The top scale corresponds to the time in millions of years.

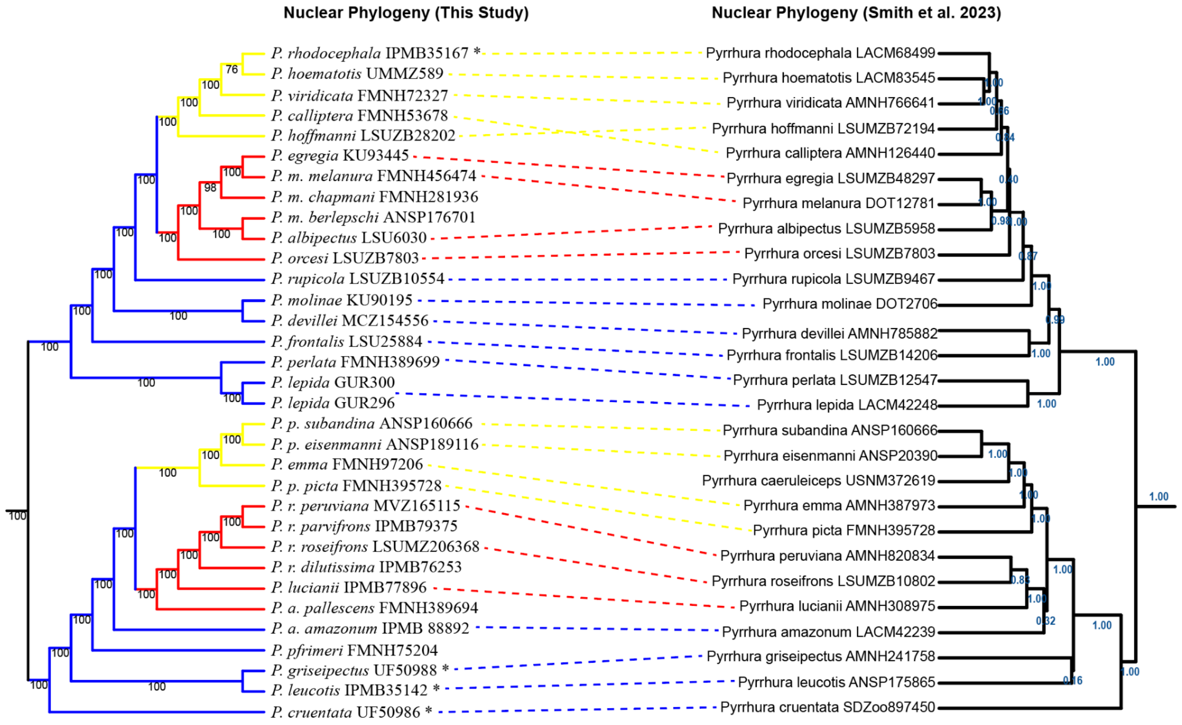

**Figure S3.** Topological comparison between the species tree estimated by Smith et al. (2023) and the species tree inferred in this study. Both analyses were performed using ASTRAL. The Smith et al. (2023) topology is based on UCE data, with posterior probabilities shown in blue. The topology generated in this study is based on 25 kb windows derived from whole-genome sequence data, and node labels represent multilocus bootstrap values.

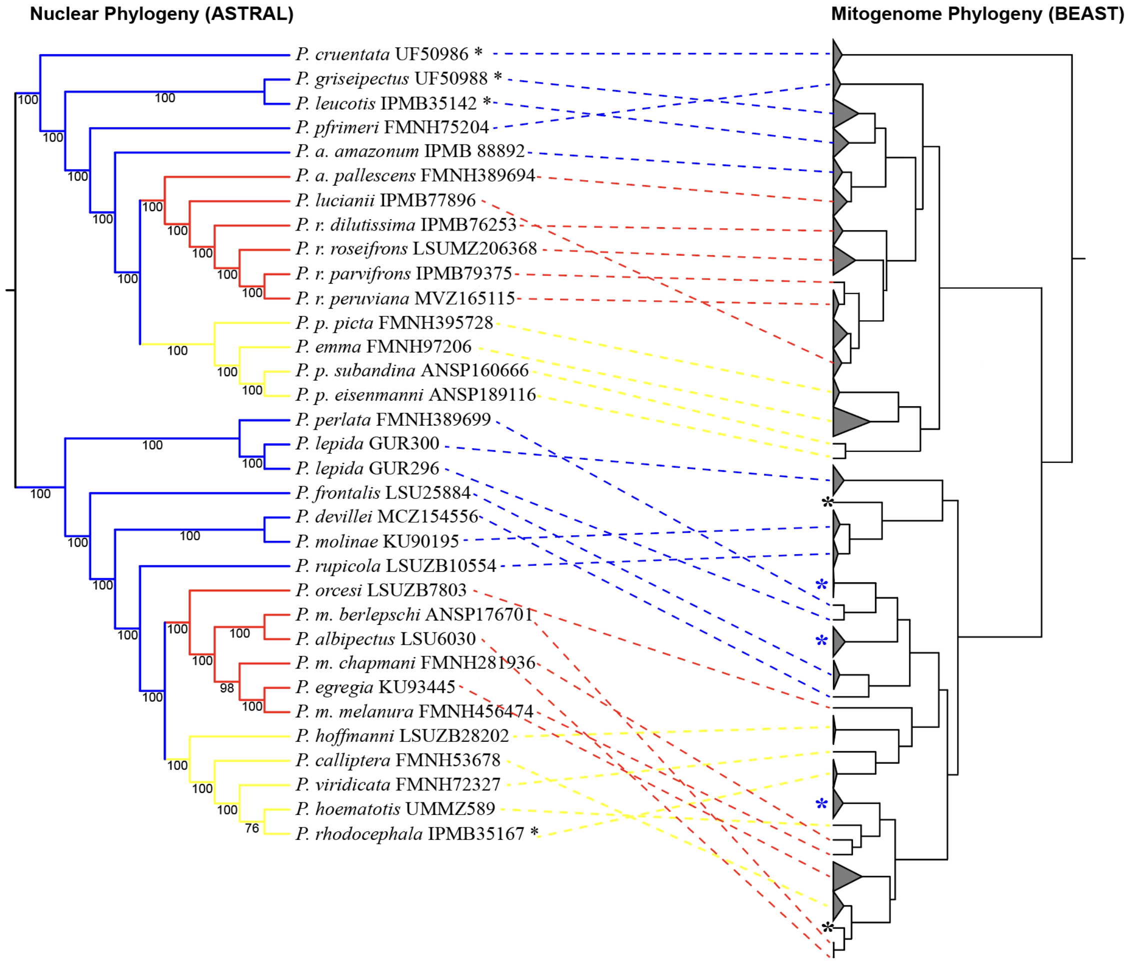

**Figure S4.** Topology discordances between this study's nuclear-based phylogeny inferred using ASTRAL (left) and mitochondrial-based phylogeny estimated using BEAST 2.7 (right). Captive clades are shown in blue asterisk, and divergent captive samples are shown in black asteriks.

**Figure S5.** Mitogenome sequence of *P. parvifrons* (MUSM 32474) and visualization of the annotation results. The inner circle shows the MITOS2 annotations, and the following corresponds to the tRNA scan results. The two upper rings correspond to the consensus annotation used to annotate the rest of the mitochondrial genomes. Mitogenome visualized in Geneious Prime 2025.1.1

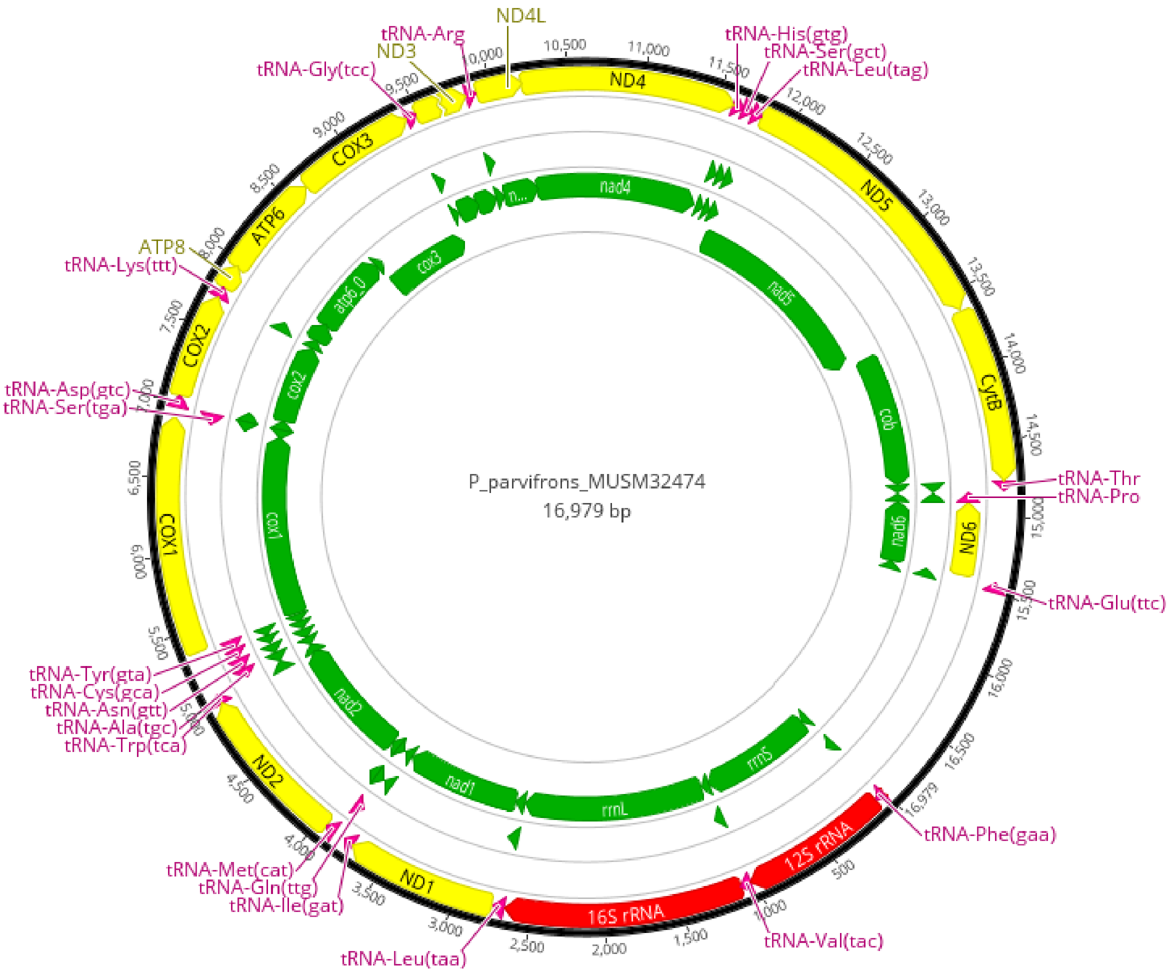
